# Mapping protein-DNA interactions with DiMeLo-seq

**DOI:** 10.1101/2022.07.03.498618

**Authors:** Annie Maslan, Nicolas Altemose, Reet Mishra, Jeremy Marcus, Lucy D. Brennan, Kousik Sundararajan, Gary Karpen, Aaron F. Straight, Aaron Streets

**Affiliations:** Department of Bioengineering, University of California, Berkeley, CA 94720; UC Berkeley-UCSF Graduate Program in Bioengineering, University of California, Berkeley, Berkeley, CA 94720; Center for Computational Biology, University of California, Berkeley, CA 94720; Department of Molecular & Cell Biology, University of California, Berkeley, CA 94720; Department of Biochemistry, Stanford University, Stanford, CA 94305; Chan Zuckerberg Biohub, San Francisco, CA 94158

## Abstract

We recently developed **Di**rected **Me**thylation with **Lo**ng-read **seq**uencing (DiMeLo-seq) to map protein-DNA interactions genome wide. DiMeLo-seq is capable of mapping multiple interaction sites on single DNA molecules, profiling protein binding in the context of endogenous DNA methylation, identifying haplotype specific protein-DNA interactions, and mapping protein-DNA interactions in repetitive regions of the genome that are difficult to study with short-read methods. With DiMeLo-seq, adenines in the vicinity of a protein of interest are methylated in situ by tethering the Hia5 methyltransferase to an antibody using protein A. Protein-DNA interactions are then detected by direct readout of adenine methylation with long-read, single-molecule, DNA sequencing platforms such as Nanopore sequencing. Here, we present a detailed protocol and practical guidance for performing DiMeLo-seq. This protocol can be run on nuclei from fresh, lightly fixed, or frozen cells. The protocol requires 1-2 days for performing in situ targeted methylation, 1-5 days for library preparation depending on desired fragment length, and 1-3 days for Nanopore sequencing depending on desired sequencing depth. The protocol requires basic molecular biology skills and equipment, as well as access to a Nanopore sequencer. We also provide a Python package, *dimelo*, for analysis of DiMeLo-seq data.

**Key papers:** Altemose, N., Maslan, A., Smith, O.K., Sundararajan, K., Brown, R.R., Mishra, R., Detweiler, A.M., Neff, N., Miga, K.H., Straight, A.F. and Streets, A., 2022. DiMeLo-seq: a long-read, single-molecule method for mapping protein–DNA interactions genome wide. *Nature Methods*, pp.1-13. (https://doi.org/10.1038/s41592-022-01475-6)

## Introduction

Common methods for mapping protein-DNA interactions rely on selective amplification and sequencing of short DNA fragments from regions bound by the protein of interest^1–7^. These powerful short-read methods for profiling protein-DNA interactions have been used to map the binding patterns of thousands of proteins in human cells^8^. However, because the measurement uses amplified fragments of DNA, these methods dissociate joint binding information at neighboring sites, remove endogenous DNA methylation, and are limited in detecting haplotype-specific interactions and interactions in repetitive DNA. DiMeLo-seq addresses these limitations by using long-read, single-molecule sequencing to measure protein-DNA interactions genome-wide (Figure 1)^9^.

**Figure 1.**
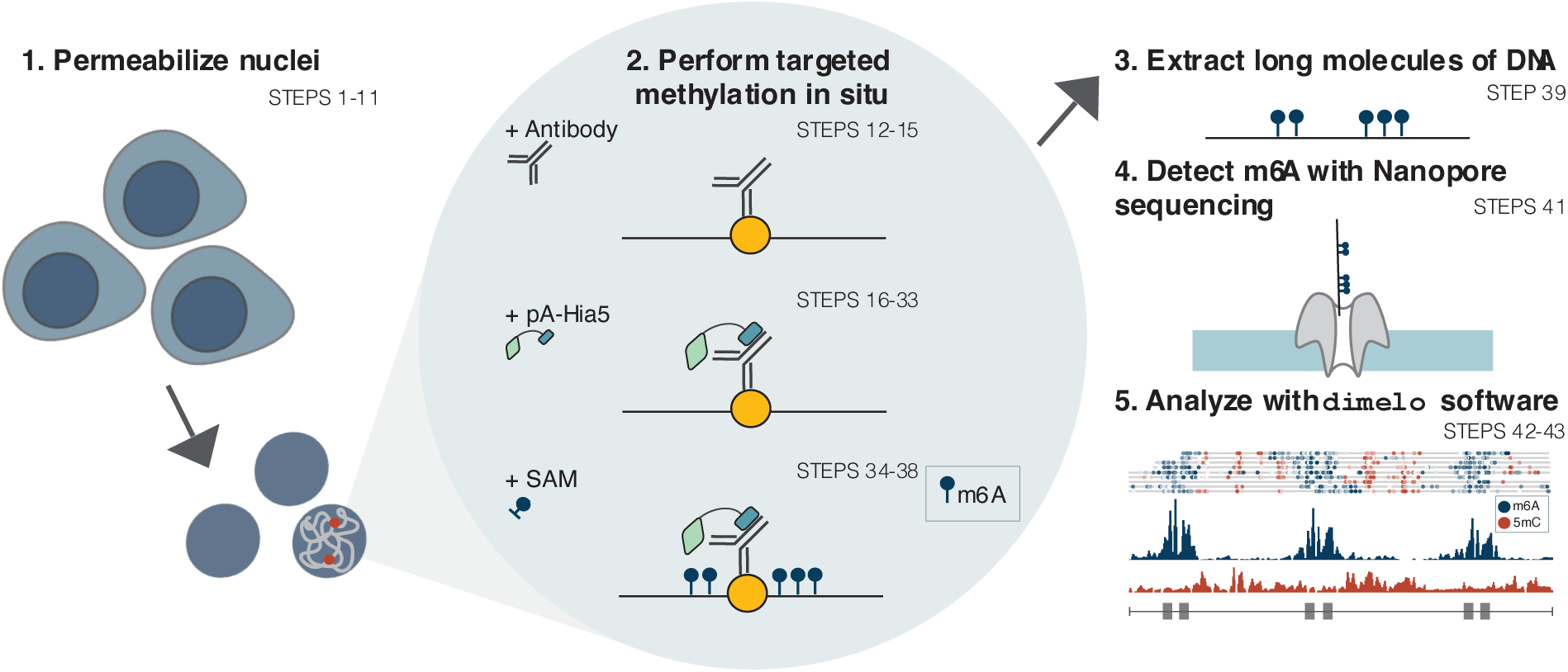
DiMeLo-seq protocol overview. Step numbers in the procedure are indicated. 1. Permeabilize nuclei from fresh, frozen, or fixed cells. 2. Perform a series of steps within the permeabilized nuclei: (i) bind primary antibody to the protein of interest, (ii) bind pA-Hia5 to the primary antibody, (iii) add S-adenosylmethionine (SAM), the methyl donor, to activate methylation. 3. Extract long molecules of DNA. 4. Sequence this DNA with a Nanopore sequencer to detect m6A directly. 5. Analyze modified basecalls from sequencing using the *dimelo* software package.

DiMeLo-seq relies on antibody-targeted DNA methylation to record protein binding events on genomic DNA in situ, followed by direct readout of methylation with single-molecule sequencing of native DNA. Single-molecule sequencing technologies, such as Oxford Nanopore Technologies (ONT) sequencers and the Pacific Biosciences Sequel IIe, can directly detect DNA modifications, including endogenous CpG methylation and exogenous adenine methylation deposited by DiMeLo-seq to mark protein binding events. Sequencing single, native DNA molecules, as opposed to sequencing short, amplified clones, enables simultaneous detection of endogenous CpG methylation and protein binding on the same read. Additionally, these single-molecule sequencing platforms can sequence much longer reads compared to those generated with Illumina sequencing, producing reads up to a megabase in length and enabling detection of multiple protein binding events across long distances. Furthermore, long-read, single-molecule sequencing makes DiMeLo-seq capable of detecting protein binding events in a truly genome-wide fashion, including repetitive regions of the genome where short reads cannot be accurately positioned.

### Development of the protocol

DiMeLo-seq was inspired by the targeted methylation strategy used in DamID-seq^7^ and builds from short-read techniques for mapping protein-DNA interactions (e.g. CUT&Tag^5^, CUT&RUN^6^, and pA-DamID^10^), as well as recent work that implements long-read sequencing and detection of exogenous methylation to profile chromatin accessibility (Fiber-seq^11^, SMAC-seq^12^, SAMOSA^13^, NanoNOMe^14^, MeSMLR-seq^15^). Developing DiMeLo-seq required substantial optimization, with over 100 conditions tested^9^. These optimization experiments revealed critical protocol components that improved efficiency including these key observations: (1) The methyltransferase Hia5 performs significantly better than EcoGII in situ; (2) digitonin and Tween-20 dramatically increase methylation levels compared to other detergents for nuclear permeabilization; and (3) the following protocol parameters all improve methylation levels: a low salt concentration, inclusion of BSA in all buffers, increased incubation times, and replenishment of the methyl donor during activation.

### Applications of the method

DiMeLo-seq can be used to profile the genome localization of any DNA-binding protein for which there is a specific, high-quality antibody. The protocol can be used on fresh, fixed, or frozen cells from culture or from primary tissue. Because DiMeLo-seq uses antibody-based targeting, DiMeLo-seq is also able to profile post-translational modifications like protein acetylation, methylation, and phosphorylation. In our previous work^9^, we demonstrated application of DiMeLo-seq for profiling binding landscapes of LMNB1, CTCF, H3K9me3, and CENPA in cultured human cells. Here, we applied DiMeLo-seq to profile H3K27ac, H3K27me3, and H3K4me3 in cultured human cells and H3K9me3 in *D. melanogaster* embryos.

### Comparison with other methods

The key distinguishing feature of DiMeLo-seq compared to other methods is that protein-DNA interactions are measured on long, native, single molecules of DNA. Long reads facilitate mapping of protein-DNA interactions and histone modifications in highly repetitive regions of the genome, measurement of multiple binding events on the same chromatin fiber, and detection of haplotype-specific interactions. Measurement of native DNA molecules allows for binding assessment in the context of endogenous CpG methylation. Additionally, single-molecule sensitivity enables measurement of heterogeneous binding at a given locus.

Long-read sequencing has enabled the recent development of a handful of new methods for profiling chromatin. For example, Fiber-seq^11^, SMAC-seq^12^, SAMOSA^13^, NanoNOMe^14^, and MeSMLR-seq^15^ are all methods that profile chromatin accessibility on long molecules of DNA and can infer protein-DNA binding footprints, although they do not directly measure specific target proteins. Furthermore, recent studies described BIND&MODIFY^16^ and nanoHiMe-seq^17^, methods which use similar strategies as DiMeLo-seq to profile protein-DNA interactions, albeit with lower reported sensitivity.

Traditional methods for profiling protein-DNA interactions such as ChIP-seq, CUT&RUN, CUT&Tag, and DamID-seq all rely on amplification of short DNA fragments, producing sequencing reads that are hundreds of base pairs in length. These methods use coverage as a proxy for binding and exhibit an inverse relationship between resolution and read length. Short reads often preclude haplotype phasing, mapping to repetitive regions, and measuring multiple binding events on the same DNA molecule. Additionally, these short-read methods require amplification, thereby losing the endogenous mCpG marks. Joint protein binding and mCpG measurement can be done with BisChIP-Seq/ChIP-BS-Seq, but this requires lossy and harsh bisulfite conversion that degrades DNA^18,19^. ChIP-seq also requires physical separation of protein-bound DNA regions and extensive washes, which can reduce sensitivity and prohibit single-cell resolution. DiMeLo-seq is a complimentary method that provides a solution to the limitations listed above. Moreover, like other antibody-based approaches, such as ChIP-seq, CUT&Tag, and CUT&RUN, DiMeLo-seq is compatible with primary cells and can be used to target post-translational modifications.

DamID-seq is a method that reports protein-DNA interactions without use of antibodies, instead relying on a fusion of an exogenous methyltransferase (Dam) and a protein of interest to methylate DNA in vivo. The methylated sites are then digested with a methylation-sensitive restriction enzyme to convert methyl marks into a signal that can be detected with Illumina sequencing. Dam is an adenine methyltransferase which recognizes GATC sites, limiting the resolution of this method in GATC-depleted regions of the genome. DamID-seq requires a genetically tractable system for expression of the Dam fusion protein. Further, optimizing the Dam fusion expression levels and induction times present major hurdles in adopting this method compared to antibody-based approaches. Importantly, with DamID-seq, binding patterns of the introduced Dam fusion rather than the endogenous protein are measured. Thus, DamID-seq is a powerful approach for profiling protein-DNA interactions and for capturing an integrated signal of where a protein has bound over the induction time period, but these limitations have made DamID-seq adoption slower relative to antibody-based approaches. We have demonstrated that cells expressing a methyltransferase fusion construct can also be sequenced with long-read platforms, combining the benefits of DamID-seq and DiMeLo-seq^9^.

Because there is no amplification step, DiMeLo-seq experiments require substantial input (typically ∼1 million cells) to generate sufficient material for sequencing. With short-read methods such as CUT&Tag and DamID-seq, protein-bound regions are selectively amplified and sequenced, thereby enriching for protein-bound regions and allowing for input as low as a single cell. While DiMeLo-seq cannot provide genome-wide coverage of single cells, the single-molecule sensitivity of long-read sequencing effectively provides single-cell resolution at a given genomic site, allowing the user to measure cell-to-cell heterogeneity of protein binding at that site. We assessed the effective single-cell resolution of DiMeLo-seq in a previous study^9^ showing a strong correlation between DiMeLo-seq interaction frequency and single-cell DamID-seq interaction frequency. Without enrichment for regions of interest, DiMeLo-seq will sequence the whole genome uniformly, requiring deep sequencing to achieve sufficient coverage of specific target regions. However, there are options to enrich for regions of interest with DiMeLo-seq like AlphaHOR-RES^9^, other restriction-enzyme-based enrichment approaches^20^, and the Oxford Nanopore Technologies Cas9-based targeted library preparation kit (SQK-CS9109). Adaptive sampling methods could also be used, such as Readfish^21^, UNCALLED^22^, and the adaptive sampling that is built into ONT’s MinKNOW software.

### Experimental design

#### Cell preparation

We have applied DiMeLo-seq to profile protein-DNA interactions in human cell lines, including GM12878, HEK293T, HG002, and Hap1, and in *D. melanogaster* embryos here and in our previous study.^9^ Cell lines are cultured under the recommended conditions and are harvested for DiMeLo-seq typically at 75-100% confluency. To harvest the cells, we perform a single wash with cold PBS and then begin nuclear permeabilization. For details of *D. melanogaster* embryo preparation, refer to Supplementary Methods.

#### Cell-type-specific and target-specific experimental parameters

While the standard DiMeLo-seq protocol described here has performed consistently for all cell types tested, application to other cell types and primary tissue may require tuning of digitonin concentration for efficient nuclear permeabilization. A digitonin concentration of 0.02% (wt/vol) has worked well for human GM12878, HG002, Hap1, HEK293T, and Drosophila embryos. The optimal digitonin concentration can be determined using Trypan Blue (Figure 2a-b). Successful permeabilization allows Trypan Blue to localize to the nucleus (Figure 2b). If too little digitonin is used, Trypan blue internalization is sparse, as in Figure 2a. If too much digitonin is used, fewer nuclei are recovered after digitonin treatment relative to the number of cells input. Recovery around 80-90% should be expected and all nuclei should appear permeabilized as in Figure 2b. Primary tissue also requires upstream processing for nuclear extraction before the nuclear permeabilization step (Supplementary Methods). It is important that the nuclear extraction method does not contain NP-40, as we have found this detergent can significantly reduce methylation. As long as a sufficient number of nuclei can be collected, DiMeLo-seq can be applied to any cell type or primary tissue. We typically start with 1 to 5 million cells for a single protein target. We do not see appreciable loss of nuclei during the nuclear permeabilization step for the cell types we have tested with 0.02% (wt/vol) digitonin, so it is sufficient to quantify input cell count rather than quantifying nuclei after permeabilization. However, with other cell types and detergent concentrations, nuclei loss during permeabilization may become more significant. In these cases, counting nuclei after permeabilization may be advisable to assure there are 1 to 5 million nuclei for the primary antibody binding step. When preparing cells for multiple experiments or targets, we often permeabilize nuclei in a single batch; we typically start with up to 20 million cells in 1 ml of Dig-Wash, divide nuclei into separate tubes for each target, and then continue with primary antibody binding for each experiment in parallel.

**Figure 2.**
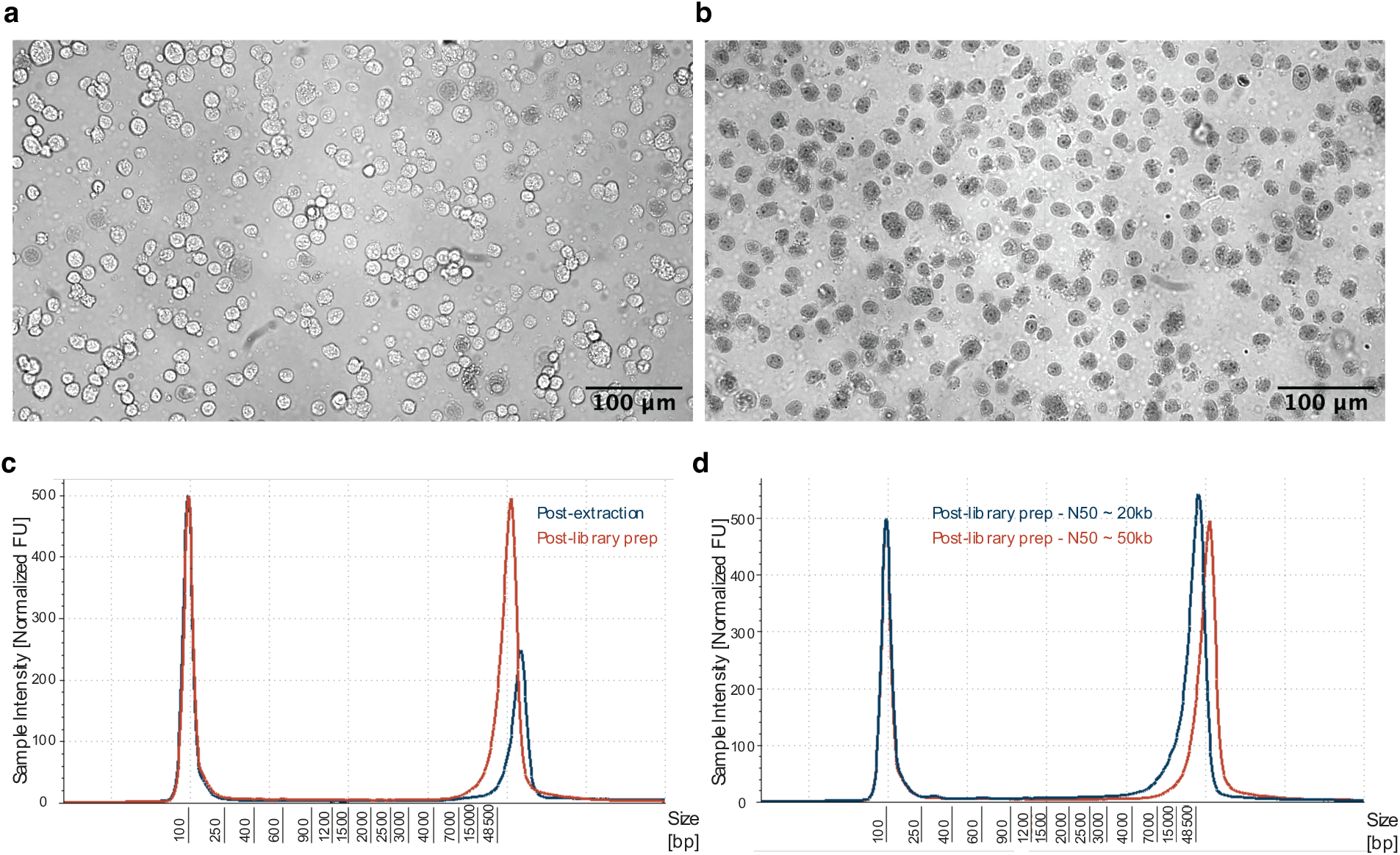
Experimental quality control. **a-b,** To determine successful permeabilization, cells are stained with Trypan Blue before (a) and after (b) digitonin treatment. Successful permeabilization allows Trypan Blue to enter the nuclei, while still maintaining high recovery of nuclei from cells. Over-permeabilization results in lower recovery of nuclei. Under-permeabilization does not allow Trypan Blue to enter the nuclei. **c-d,** After the DiMeLo-seq in situ protocol and DNA extraction, DNA is sized using the TapeStation. Representative traces from ligation-based library preparation are show in (c) for the fragment size distribution after extraction and after library preparation. In (d), the size distribution after library preparation for the two ligation-based methods presented in this protocol are shown. The blue curve results in N50 ∼20 kb, while the red curve results in N50 ∼50 kb. Larger fragment sizes can be achieved with other ultra long kits.

When working with new protein targets, key variables to optimize are the antibody concentration and the extent of fixation for targets with low binding affinity. An antibody dilution of 1:50 has worked well for all targets reported here and in Altemose et al.^9^ Extensive washes are performed after antibody binding to remove any unbound antibody, making excess antibody less detrimental. We have demonstrated that light fixation is compatible with the DiMeLo-seq workflow. If targeting a protein that binds transiently, including fixation may improve signal by preventing the protein from dissociating from the DNA during the DiMeLo-seq protocol. However, care should be taken to evaluate the susceptibility of any new protein targets to epitope masking by fixation^23^.

#### Recommended controls

Typical controls include an IgG isotype control and a free pA-Hia5 control. The IgG isotype control measures nonspecific antibody binding. The free pA-Hia5 control measures chromatin accessibility, similar to Fiber-seq and related methods, and is analogous to the Dam only control used in DamID-seq^11,24^. This control is performed by excluding the primary antibody and pA-Hia5 binding steps and instead adding pA-Hia5 at activation at 200 nM. While these controls are not required, they provide a useful measure of background methylation and bias caused by variable chromatin accessibility. Excluding pA-Hia5 as a control to account for modified basecalling errors can also be included. If troubleshooting a DiMeLo-seq experiment, using one of the antibodies and cell lines validated here and in Altemose et al.^9^ may also be a useful control.

#### Choice of commercial kits and sequencing platforms

Commercially available kits and techniques for DNA extraction, library preparation, and sequencing are rapidly improving. The protocol described here produces consistent localization profiles shown below and in Altemose et al.^9^, but it is important to note that after the in situ methylation steps, any DNA extraction method, library preparation kit, flow cell chemistry, and sequencing device can be used as long as m6A is maintained (no amplification is performed) and the flow cell and device have basecalling models available for calling m6A. We have also demonstrated sequencing of DiMeLo-seq samples with Pacific Biosciences’ Sequel IIe^9^.

#### Sequencing considerations

The key considerations for sequencing are fragment size and sequencing depth. The target N50 (half of sequenced bases are from a fragment size of N50 or larger) varies by application. Longer reads may be desired when mapping to repetitive regions, probing coordinated binding events at longer distances, or phasing reads. For example, if interested in studying multiple binding events on single molecules, the target protein’s binding site density and the number of sites the user would like to capture on a single molecule should inform the desired fragment size. With ligation-based library preparation we typically target an N50 of ∼20 kb to ∼50 kb, which results in fragment size distributions as in Figure 2c-d. With other library preparation kits (e.g., SQK-ULK114), much larger fragments can be sequenced; however, there is a tradeoff between fragment length and throughput. If a user targets larger fragment sizes, the total sequencing output for the flow cell, and thus total coverage, will be reduced. Greater sequencing depth provides more accurate and sensitive recovery of binding landscapes. This target sequencing depth will also vary by application, depending on the binding footprint of the protein, the mappability of the region of interest, and the biological question at hand. Final libraries can be saved, and flow cells can be reloaded, so it is recommended to do an initial pilot run with shallow sequencing followed by deeper sequencing as needed. For example, for initial tests of new protein targets in human cells, we typically sequence to 0.3-1X coverage (∼1-3 Gb) to validate and determine optimal experimental conditions, and then sequence more deeply to ∼5-45X coverage depending on the analysis we are performing. For example, when targeting CTCF, we sequenced to ∼25X coverage and detected 60% of ChIP-seq peaks with a false positive rate of 1.6%^9^. See the sequencing saturation analysis with CTCF-targeted DiMeLo-seq in Altemose et al.^9^

#### Additional experimental considerations

The protocol will be kept up-to-date at https://www.protocols.io/view/dimelo-seq-directed-methylation-with-long-read-seq-n2bvjxe4wlk5. The version used for this manuscript is Version 2 (https://doi.org/10.17504/protocols.io.b2u8qezw). Sequencing a single sample with the product list described in this manuscript costs roughly $1100. This calculation assumes an average antibody price and ligation-based library preparation without multiplexing. On a MinION flow cell, this yields ∼25 Gb data, or ∼8X coverage of the human genome. On a PromethION flow cell, this yields ∼125 Gb data, or ∼40X coverage of the human genome.

Sequencing a single sample on a MinION flow cell yielding ∼25 Gb data, or ∼8X coverage of the human genome, costs roughly $1100 with the product list described in this manuscript. This calculation assumes an average antibody price and ligation-based library preparation without multiplexing.

All spins are at 4°C for 3 minutes at 500 x g. Spinning in a swinging bucket rotor can help pellet the nuclei. To prevent nuclei from lining the side of the tube, break all spins into two parts: 2 minutes with the tube hinge facing inward, followed by 1 minute with the tube hinge facing outward. This two-part spin is not needed if using a swinging bucket rotor. Working with Eppendorf DNA LoBind tubes can reduce loss of material. Use wide bore tips when working with nuclei. Do not use NP-40 or Triton-X100 for nuclear extraction, permeabilization, or any other stage of the protocol, as they appear to dramatically reduce methylation activity. We use Tween-20 to reduce hydrophilic non-specific interactions and BSA to reduce hydrophobic non-specific interactions. We also found that including BSA at the activation step significantly increases methylation activity. For pA-incompatible antibodies, a secondary antibody can be used as a bridging antibody, but performance is diminished; instead, we recommend using pA/G-Hia5 for pA-incompatible antibodies. See Supplementary Table 1 for a performance comparison for pA/G-Hia5 and pA-Hia5. While both achieve a comparable on-target to off-target methylation ratio, pA/G-Hia5 on-target methylation rates are slightly lower that those with pA-Hia5 when targeting LMNB1 in HEK293T cells.

We have found that the pA-Hia5 fusion protein can be stored at -80°C for at least 6 months without decline in performance. We typically spin down the pA-Hia5 protein aliquots at 10,000 x g for 10 minutes at 4°C before each experiment to remove insoluble protein and to confirm that there has been no reduction in soluble protein concentration.

#### Analysis

We have created a Python package called *dimelo*^25^ for analysis of DiMeLo-seq data (https://github.com/streetslab/dimelo). The *dimelo* package input format requires raw output files produced by the Nanopore sequencer to first be converted to BAM files. Recommendations for the basecalling and alignment steps, which will create an aligned BAM file with “Mm” and “Ml” tags that describe methylation calls, can be found in the package documentation (https://streetslab.github.io/dimelo/). For this manuscript, we have used Megalodon (v 2.3.1) and Guppy (v4.5.4) with the Rerio res_dna_r941_min_modbases-all-context_v001.cfg basecalling model. An ONT account is useful for accessing user manuals and for troubleshooting. Basecalling is being rapidly improved by ONT and others, so basecalling suggestions reported here are likely to become outdated quickly. After basecalling and alignment, the resulting BAM file is the input to the quality control, visualization, and custom analysis functions from the *dimelo* software package. This analysis software can be run as an imported Python module or from the command line. Figure 3 provides a summary of the functions included in the *dimelo* package.

**Figure 3.**
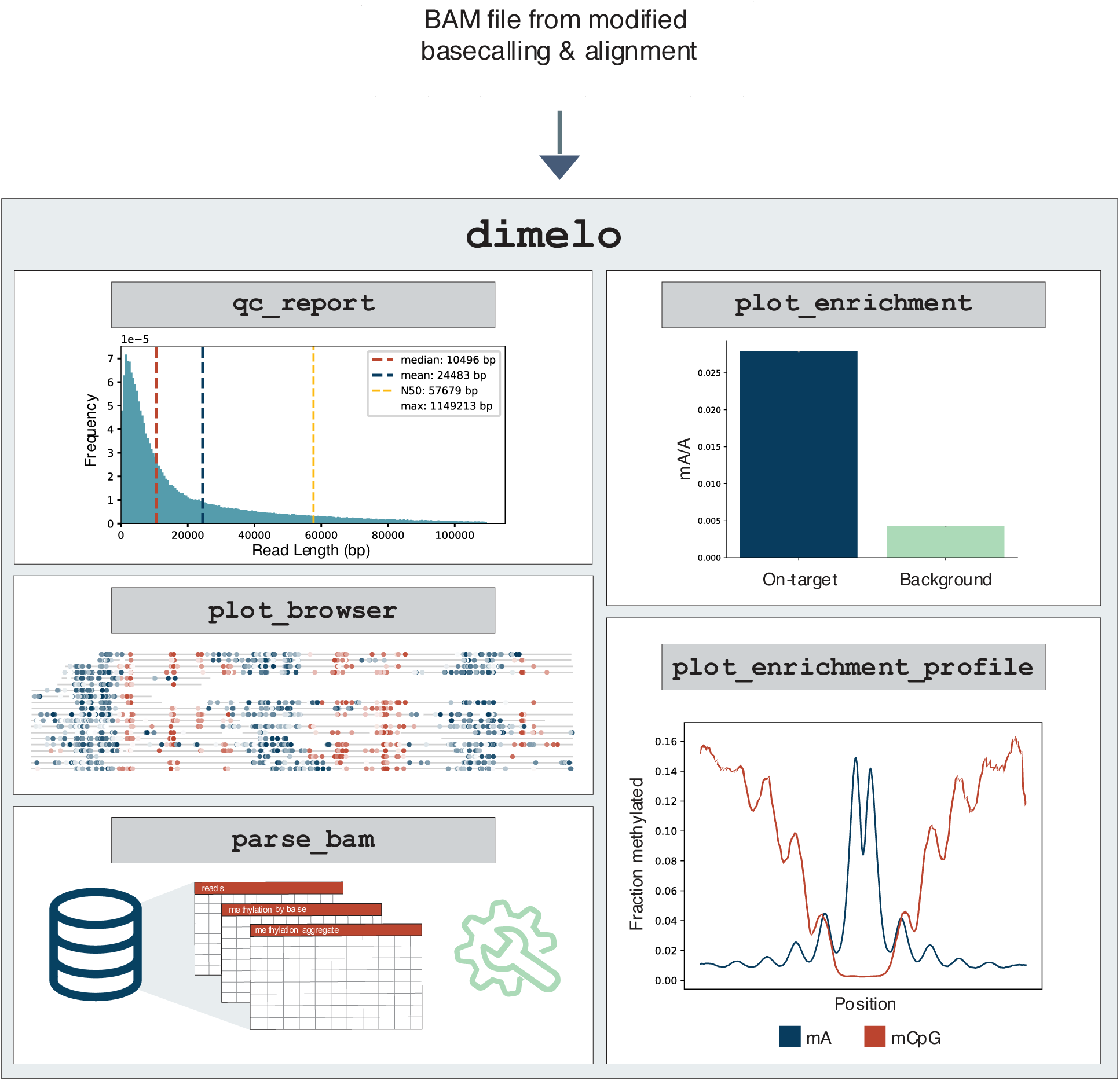
Analysis pipeline overview. Basecalling and alignment are performed on the FAST5 output from the Nanopore sequencer. The resulting BAM that contains the modified base information is then input to the *dimelo* software package. A recommended workflow involves quality control with `qc_report`, followed by visualization with `plot_browser`, `plot_enrichment`, and `plot_enrichment_profile`. For custom analysis, `parse_bam` stores base modification calls in an intermediate format that makes it easier to manipulate for downstream analysis.

A recommended workflow is to first run `qc_report` to generate summary statistics and histograms for metrics such as coverage, read length, mapping quality, basecall quality, and alignment quality (Figure 4). Next, three functions are provided for visualization: `plot_enrichment`, `plot_browser`, and `plot_enrichment_profile`. All functions take BAM file(s) as input and region(s) of interest defined as a string or BED file.

**Figure 4.**
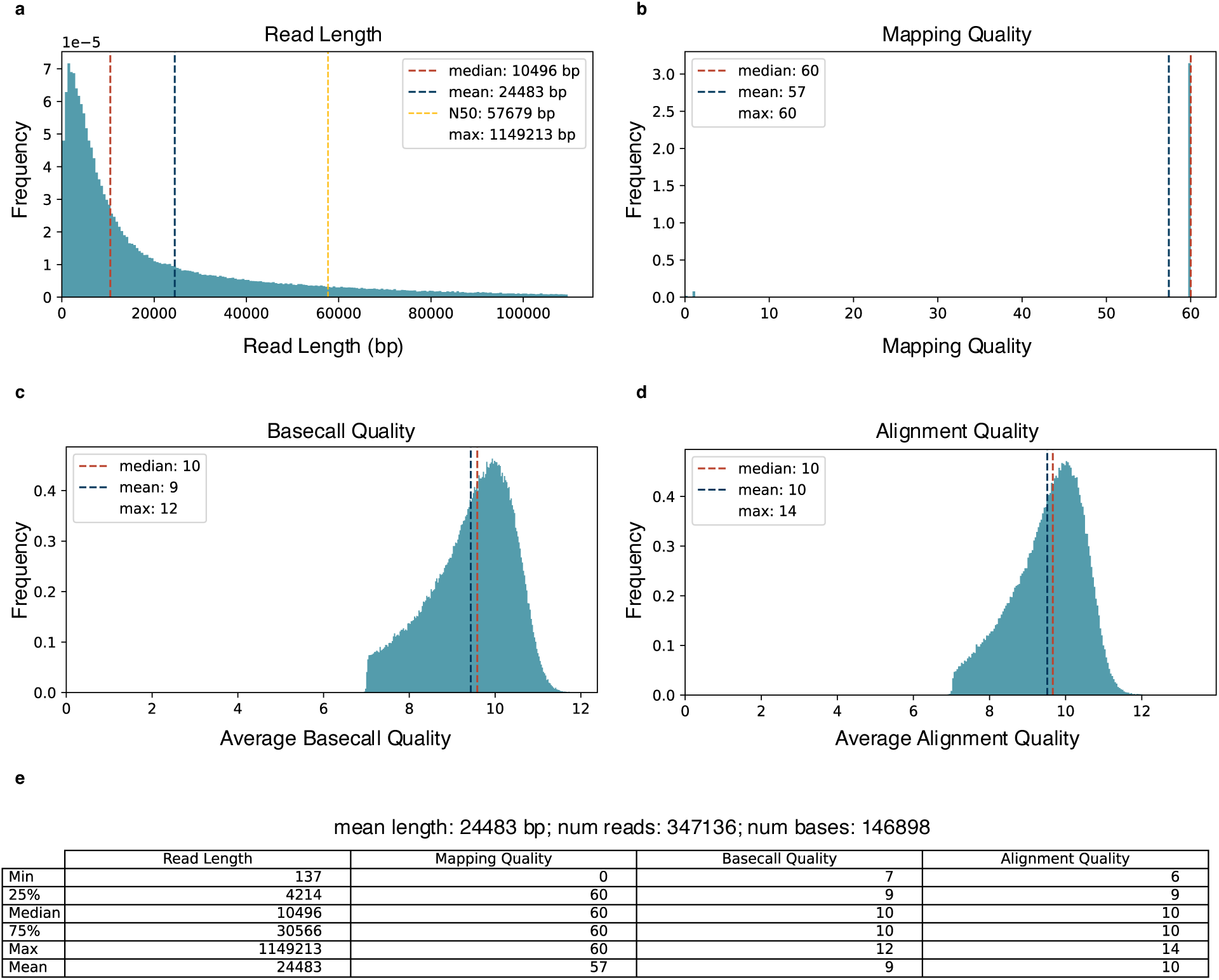
Sequencing quality control. The `qc_report` function takes in one or more BAM files and for each, outputs a QC report including the following 5 features. **a,** Histogram of read lengths with the median, mean, N50, and max value annotated. **b,** Histogram of mapping quality. **c-d,** Both the basecall quality and alignment quality scores are present in BAM outputs from Guppy but not from Megalodon. **c,** Histogram of average basecall quality per read. Here, the mean indicates that our sample’s average basecall quality is Q10, which is equivalent to 90% accuracy. **d,** Histogram of average alignment quality per read. While mapping quality provides the accuracy of the read mapping to specific genomic coordinates, the average alignment quality provides the quality of matching between the read and the reference sequence. For example, if a read almost perfectly matches multiple genomic coordinates, it will have a low mapping quality but a high alignment quality. **e,** Summary table with descriptive statistics of each feature (a-d), in addition to highlighting important values such as mean length of reads, total number of reads, and total number of bases sequenced. Example data used in this figure are from targeting H3K9me3 in *D. melanogaster* embryos.

The `plot_enrichment` function compares methylation levels across samples or across different genomic regions. This tool is useful for looking at overall on- vs. off-target methylation and for comparing methylation levels in regions of interest across samples.

The `plot_browser` function allows the user to view single molecules with base modifications colored according to the probability of methylation within a region of interest. This function can either produce a static PDF of the single molecules or an interactive HTML file that allows the user to zoom and pan around the browser plot, using plotting code adapted from Methplotlib.^26^ Plots of aggregate coverage and the fraction of methylated bases over the window of interest are also generated with this function.

The `plot_enrichment_profile` function creates single-molecule plots and an aggregate plot of the fraction of methylated bases centered at features of interest defined in a BED file. For example, one may enter a BED file with the locations of features of interest (i.e. the binding motif for a given protein, transcription start sites, etc.) to view the methylation profiles around those features. Inputting multiple BAM files creates an overlay of the methylation profiles across samples, while inputting multiple BED files creates an overlay of methylation profiles for a given sample across the different sets of regions defined in the BED files.

The `parse_bam` function summarizes the base modification information from a BAM file and stores it into an easy to manipulate data format. This gives users the ability to more easily create custom figures and analyses.

For all functions, the user can specify the modification(s) of interest to extract: “A”, “CG”, or “A+CG”. The probability threshold for calling a base as modified is also a parameter to each function. For discussion of threshold determination see Supplementary Note 6 of Altemose et al.^9^

### Expertise needed to implement the protocol

To perform DiMeLo-seq and analyze the data produced, basic molecular biology skills and basic command line skills are required. Experience working with long molecules of DNA is also beneficial, as care must be taken to maintain long fragments for sequencing^27^.

### Limitations

The performance of DiMeLo-seq is strongly dependent on the quality of the antibody used to target the protein of interest. For proteins that do not have a specific, high-quality antibody compatible with protein A, one could consider epitope tagging or performing in vivo expression of a protein-methyltransferase fusion^28^. The activity of pA-Hia5 can vary across enzyme preparations; if performing comparative studies, it may be advisable to perform all experiments with the same enzyme batch. Enzyme activity can also be evaluated before use, as described in the Supplementary Methods. DNA must be accessible for Hia5 to efficiently methylate in situ. Thus, targets in less accessible regions of the genome may require longer incubations or deeper sequencing. Additionally, this preferential methylation of open chromatin results in signal bias. Hia5 may also methylate DNA in trans if three-dimensional contacts bring topologically associated loci close enough to the target protein, potentially creating non-specific background. Such artifacts should be accounted for if absolute measurement of binding frequency is desired. DiMeLo-seq experiments typically require ∼1 million cells as input, although Concanavalin A beads (which we previously showed are compatible with DiMeLo-seq) and lower-input library preparation kits can reduce required input material^9^. Unlike other protein-DNA interaction mapping methods that enrich protein-bound DNA, the standard DiMeLo-seq protocol and sequencing library preparation yields uniform coverage of the entire genome. If only particular regions of the genome are of interest, the protocol can be modified with an enrichment step or targeted sequencing library preparation.

The resolution of DiMeLo-seq can be considered both at the single molecule level and in ensemble measurements. This resolution is dependent on the choice of target and antibody, the adenine density of the target locus, and the local chromatin environment. Across molecules in aggregate, we have observed a slightly larger binding footprint with DiMeLo-seq compared to other methods. For example, when targeting CTCF, DiMelo-seq measures an 88 bp footprint compared to ∼50 bp for methods like DNase I footprinting and ChIP-exo. This is likely because Hia5 cannot reach the DNA directly adjacent to the binding event within ∼20 bp^9^. On single molecules, DiMeLo-seq can localize the center of binding footprints to within ∼200 bp; however, this may be improved with optimized peak calling algorithms. Single-molecule resolution is also related to the measurement sensitivity, which can vary according to the same parameters as resolution. Therefore, both sensitivity and resolution are reduced in heterochromatin and in GC-rich regions of the genome.

Here and in our previous study, we have benchmarked DiMeLo-seq’s performance in targeting LMNB1, CTCF, H3K9me3, CENPA, H3K27ac, H3K27me3, and H3K4me3^9^. When targeting CTCF and LMNB1, we estimated 54% and 59% sensitivity (94% specificity), but this is dependent on the protein, antibody, and chromatin environment and must be evaluated for new targets.^9^ Transiently bound proteins may benefit from the optional fixation step at the start of the DiMeLo-seq protocol; however, as described by others^29^, we have observed that fixation decreases DNA fragment sizes. High levels of fixation are also known to mask epitopes and prevent antibody binding^23^. Therefore, the user should weigh the possible benefits of fixation against the potential fragment size hit at sequencing and potential reduction in signal. We developed a simple peak calling algorithm to identify binding sites de novo and used this peak caller to identify CTCF binding sites. At ∼25X coverage, 60% of ChIP-seq peaks are detected with a false positive rate of 1.6%^9^. In our previous study, we found that among the peaks detected with DiMeLo-seq that were not annotated ChIP–seq peaks, 10% overlapped 1-kb marker deserts and gaps in the hg38 reference and were undetectable by ChIP–seq. Another 12% of these peaks fell within 500 bp of a known CTCF motif. Further development of peak calling algorithms which take into account the systematic biases associated with DiMeLo-seq will likely improve the sensitivity and accuracy of this technique.

## Materials

### REAGENTS

#### A. Reagents for in situ protocol

- HEPES-KOH 1 Molarity (M) pH 7.5 (Boston BioProducts BBH-75-K)
- NaCl 5 M (Sigma-Aldrich 59222C-500ML)
- Spermidine 6.4 M (Sigma-Aldrich S0266-5G)
- Roche cOmplete™ EDTA-free Protease Inhibitor Tablet (Sigma-Aldrich 11873580001)
- Bovine Serum Albumin (Sigma-Aldrich A6003-25G)
- Digitonin (Sigma-Aldrich 300410-250MG) CAUTION acute toxic and health hazard; work in fume hood when making digitonin solution.
- Tween-20 (Sigma-Aldrich P7949-100ML)
- Tris-HCl 1M pH 8.0 (Invitrogen 15568025)
- KCl (Sigma-Aldrich PX1405-1)
- EDTA 0.5 M pH 8.0 (Invitrogen 15575-038)
- EGTA 0.5 M pH 8.0 (Fisher 50-255-956)
- S-Adenosylmethionine 32 mM (NEB B9003S)
- PFA, 16% (wt/vol) (if performing fixation) (EMS 15710)
- Glycine (if performing fixation) (Fisher BP381-1)
- Eppendorf DNA LoBind tubes 1.5 mL (Fisher 022431021)
- Wide bore 200 µl and 1000 µl tips (e.g. USA Scientific 1011-8810, VWR 89049-168)
- pA-Hia5 (see https://www.protocols.io/view/pa-hia5-protein-expression-and-purification-x54v9j56mg3e for expression and purification protocol. Version 1 of the protocol is used for this manuscript (<u>dx.doi.org/10.17504/protocols.io.bv82n9ye</u>). The pET-pA–Hia5 (pA-Hia5) plasmid is available from Addgene (cat no. 174372)).
- Primary antibody for protein target of interest, from species compatible with pA (e.g. Abcam ab16048)
- Secondary antibody for immunofluorescence quality control (e.g. Abcam ab3554)
- IgG antibody - isotype control, from species compatible with pA (e.g. Abcam ab171870)
- Trypan Blue (Fisher T10282)
- Qubit dsDNA BR Assay Kit (Fisher Q32850)
- Qubit Protein Assay Kit (Fisher Q33211)

#### B. Reagents for extraction, library preparation, and sequencing

CAUTION We have validated the following reagents, but extraction, library preparation, and sequencing reagents are improving rapidly. The important considerations are to choose a DNA extraction method that maintains long DNA molecules, to perform amplification-free library preparation, and to use ONT kit chemistry that is compatible with mA calling.

- Monarch Genomic DNA Purification Kit (NEB T3010S)
- Monarch HMW DNA Extraction Kit (NEB T3050L)
- Agencourt AMPure XP beads (Beckman Coulter A63881)
- Blunt/TA Ligase Master Mix (NEB M0367S)
- NEBNext quick ligation module (NEB E6056S)
- NEBNext End Repair dA-tailing Module (NEB E7546S)
- NEBNext FFPE DNA repair kit (NEB M6630S)
- Ligation Sequencing Kit (ONT SQK-LSK109, ONT SQK-LSK110, or latest kit compatible with m6A calling)
- Native Barcoding Expansion 1-12 (ONT EXP-NBD104, or latest kit compatible with m6A calling)
- Native Barcoding Expansion 13-24 (ONT EXP-NBD114, or latest kit compatible with m6A calling)
- Circulomics Short Read Eliminator Kit (SS-100-101-01)
- Flow Cell Wash Kit (ONT EXP-WSH004)
- Flow cells (ONT FLO-MIN106D or ONT FLO-PRO002, or latest flow cells compatible with m6A calling)

### EQUIPMENT

- Centrifuge that can hold 4°C
- Rotator (e.g. Millipore Sigma Z740289) for end-over-end tube rotation
- Heat block / thermal mixer (e.g. Fisher Scientific 15-600-330)
- Oxford Nanopore Technologies Nanopore sequencer (e.g. MIN-101B)
- Magnetic separation rack (if targeting N50 ∼20 kb) (e.g. NEB S1515S)
- Qubit (e.g. Thermo Fisher Scientific Q33238)
- Tapestation (not required; for quality control) (e.g. Agilent G2992AA)
- Microscope (not required; for quality control)

### BIOLOGICAL MATERIALS

This protocol can be used for cell lines or for primary cells. In particular, we have validated the use of DiMeLo-seq in the following:

□ Cell lines (GM12878, HEK293T, HG002, and Hap1 are validated here and in Altemose et al.^9^) CAUTION The cell lines used in your research should be regularly checked to ensure they are authentic and are not infected with mycoplasma.
□ *D. melanogaster* embryos (see Supplementary Methods for processing details)

### REAGENT SETUP

#### A. Buffer preparation

Prepare all buffers fresh the day of the experiment, filter buffers through a 0.2 μm filter, and keep buffers on ice or at 4 °C.

##### Digitonin

Solubilize digitonin in preheated 95°C Milli-Q water to create a 5% (wt/vol) digitonin solution (e.g. 10 mg/200 μl).

##### Wash Buffer

Prepare wash buffer according to the following table. This is sufficient for 10 samples.

| Component | Amount | Final Concentration |
| --- | --- | --- |
| HEPES-KOH, 1 M, pH 7.5 | 1 ml | 20 mM |
| NaCl, 5 M | 1.5 ml | 150 mM |
| Spermidine, 6.4 M | 3.91 µl | 0.5 mM |
| Roche Complete tablet -EDTA | 1 tablet | - |
| BSA | 50 mg | 0.1% (wt/vol) |
| H2O (Milli-Q) | up to 50 ml | - |

##### Dig-Wash Buffer

Add 0.02% (wt/vol) digitonin to wash buffer. For example, add 20 µl of 5% (wt/vol) digitonin solution to 5 ml wash buffer. The optimal concentration of digitonin may vary by cell type.

##### Tween-Wash Buffer

Add 0.1% (vol/vol) Tween-20 to wash buffer. For example, add 50 µl Tween-20 to 50 ml wash buffer.

##### Activation Buffer

Prepare the activation buffer but wait to add SAM until the activation step. This is sufficient for 500 samples. Extra is made to avoid pipetting small volumes.

| Component | Amount | Concentration |
| --- | --- | --- |
| Tris, pH 8.0 1 M | 750 μL | 15 mM |
| NaCl 5 M | 150 μL | 15 mM |
| KCl 1 M | 3 mL | 60 mM |
| EDTA, pH 8.0 0.5 M | 100 μL | 1 mM |
| EGTA, pH 8.0 0.5 M | 50 μL | 0.5 mM |
| Spermidine, 6.4 M | 0.391 μL* | 0.05 mM |
| BSA | 50 mg | 0.1% (wt/vol) |
| H2O (Milli-Q) | up to 50 mL | - |
| SAM, 32 mM | (add at activation step) | 800 μM |

\*To reduce pipetting error, first perform a 1:10 dilution of Spermidine in H2O (Milli-Q) by adding 1 μL of 6.4 M Spermidine to 9 μL H2O (Milli-Q). Mix well and then add 3.91 μL of this dilution to the Activation Buffer.

## Procedure

### (Optional fixation)

TIMING 10 minutes

1. Resuspend cells in PBS (1 million to 5 million cells per condition).
2. Add PFA to 0.1% (wt/vol) (e.g. 6.2 µl of 16% (wt/vol) PFA to 1 ml cells) for 2 minutes with gentle pipet mixing.
3. Add 1.25 M glycine (sterile; 0.938 g in 10 ml) to twice the molar concentration of PFA to stop the crosslinking (e.g. 60 µl of 1.25 M glycine to 1 ml). Pipet mix.
4. Centrifuge 3 minutes at 500 x g at 4°C and remove the supernatant.
5. Continue with Nuclear permeabilization, starting with step 8.

### Nuclear permeabilization

TIMING 15 minutes

6. Prepare cells (1M-5M per condition).
7. Wash cells in PBS. Spin and remove supernatant.
8. Resuspend cells in 1 ml Dig-Wash buffer in a 1.5 ml DNA LoBind tube. Incubate for 5 minutes on ice. CAUTION The best digitonin concentration may vary by cell type. CRITICAL Use DNA LoBind tubes for all steps. CRITICAL Use wide bore tips when working with nuclei. CRITICAL Do not use NP-40 or Triton-X100 for nuclear extraction, permeabilization, or any other stage of the protocol, as they appear to dramatically reduce methylation activity. TROUBLESHOOTING
9. Split nuclei suspension into separate 1.5 ml DNA LoBind tubes for each condition.
10. Spin at 4°C for 3 minutes at 500 x g and remove supernatant. CAUTION To prevent nuclei from lining the side of the tube, break all spins into two parts: 2 minutes with the tube hinge facing inward, followed by 1 minute with the tube hinge facing outward. This two-part spin is not needed if using a swinging bucket rotor.
11. Quality control: Check permeabilization was successful by taking 1 µl of the nuclei following the 5-minute incubation on ice, diluting to 10 µl with PBS, and staining with Trypan Blue.

### Primary antibody binding

TIMING overnight (or 2.5 hours)

12. Gently resolve each pellet in 200 µl Tween-Wash containing primary antibody at 1:50 or the optimal dilution for your antibody and target.
13. Place on rotator at 4°C overnight. Samples can be incubated for ∼2 hr at 4°C instead of overnight if performing the in situ protocol in a single day.
14. Spin at 4°C for 3 minutes at 500 x g and remove supernatant.
15. Wash twice with 0.95 ml Tween-Wash. For each wash, gently and completely resolve the pellet. This may take pipetting up and down ∼10 times. Following resuspension, place on rotator at 4°C for 5 minutes before spinning down at 4°C for 3 minutes at 500 x g.

### Quantify pA-Hia5 concentration

TIMING 30 minutes

16. Thaw protein from -80°C at room temperature and then move to ice immediately.
17. Spin at 4°C for 10 minutes at 10,000 x g or higher to remove aggregates.
18. Transfer the supernatant to a new tube and save it, discarding the previous tube.
19. Use Qubit with 2 µl sample volume to quantify protein concentration.

### pA-Hia5 binding

TIMING 2.5 hours

20. Gently resolve pellet in 200 µl Tween-Wash containing 200 nM pA-Hia5.
21. Place on rotator at 4°C for ∼2 hr.
22. Spin at 4°C for 3 minutes at 500 x g and remove supernatant.
23. Wash twice with 0.95 ml Tween-Wash. For each wash, gently and completely resolve the pellet. Following resuspension, place on rotator at 4°C for 5 minutes before spinning down at 4°C for 3 minutes at 500 x g.

### Quality control to verify primary antibody and protein A binding (optional; only recommended for protein targets with specific nuclear localization patterns detectable by immunofluorescence (e.g., LMNB1))

TIMING 1 hour

24. Add 1.6 µl of 16% (wt/vol) PFA to 25 µl of nuclei in Tween-Wash (taken from the 0.95 ml final wash) for 1% (wt/vol) total PFA concentration.
25. Incubate at room temperature for 5 minutes.
26. Add 975 µl of Tween-Wash to stop the fixation by dilution.
27. Add 1 µl fluorophore-conjugated secondary antibody.
28. Put on rotator for 30 minutes at room temperature, protected from light.
29. Wash 2 times with Tween-Wash as in Step 23 (or just once). Pellet likely won’t be visible.
30. Resuspend in mounting media after last wash. Use as little as possible, ideally 5 µl.
31. Put 5 µl on a slide, make sure there are no bubbles, and put on a coverslip.
32. Seal with nail polish along the edges.
33. Image or put at -20°C once the nail polish has dried. TROUBLESHOOTING

### Activation

TIMING 2.5 hours

34. Gently resolve pellet in 100 µl of Activation Buffer per sample. Be sure to add SAM to a final concentration of 800 µM in the activation buffer at this step! In 100 µl of Activation Buffer, this means adding 2.5 µl of the SAM stock that is at 32 mM.
35. Incubate at 37°C for 2 hours on a heat block. Replenish SAM by adding an additional 800 µM at 1 hour. This means adding an additional 2.5 µl of the SAM stock that is at 32 mM to the 100 µl reaction. Pipet mix every 30 minutes. Tapping to mix also works.
36. Spin at 4°C for 3 minutes at 500 x g and remove supernatant.
37. Resuspend in 100 µl cold PBS.
38. Check nuclei by Trypan blue staining to determine recovery and check integrity of nuclei if desired.

### DNA extraction

39. Perform DNA extraction.

CAUTION We have validated the following reagents, but extraction, library preparation, and sequencing reagents are improving rapidly. The important considerations are to choose a DNA extraction method that maintains long DNA molecules, to perform amplification-free library preparation, and to use ONT kit chemistry that is compatible with mA calling. These are workflows we have validated and modifications we have made.

#### (A) Target N50 of ∼20 kb

TIMING 1 hour

(i) Use the Monarch Genomic DNA Purification Kit. Follow protocol for genomic DNA isolation using cell lysis buffer. Include RNase A. NB. If fixation was performed, be sure to do the 56°C incubation for lysis for 1 hour (not just 5 minutes) to reverse crosslinks.
(ii) Perform two elutions: 100 µl and then 35µl. PAUSE POINT - Samples can be stored at 4°C or -20°C.
(iii) Quantify DNA yield by Qubit dsDNA BR Assay Kit.
(iv) Concentrate by SpeedVac if necessary for 1-3 µg DNA in 48 µl for input to library prep. Do not use heat with the SpeedVac to prevent fragmenting the DNA.

#### (B) Target N50 ∼50 kb

TIMING 1 hour

(i) Use the NEB Monarch HMW DNA Extraction Kit. Follow protocol for genomic DNA isolation using cell lysis buffer. Include RNase A. Perform lysis with 2000 rpm agitation. We have validated 2000 rpm gives N50 ∼50-70 kb but if longer reads are desired we expect 300 rpm would work. Apart from using a different kit, all of the steps for the long fragment DNA extraction are the same as the general protocol. To reiterate, make the following changes to the protocol outlined in the following steps. If fixation was performed, be sure to do the 56°C incubation for lysis for 1 hour (not just 10 minutes) to reverse crosslinks. Agitate for 10 minutes and then keep at 56°C without agitation for 50 minutes. PAUSE POINT - Samples can be stored at 4°C or -20°C.
(ii) Quantify DNA yield by Qubit dsDNA BR Assay Kit.
(iii) Concentrate by speedvac if necessary to obtain 1-3 µg DNA in 48 µl for input to library prep.

### (Optional enrichment)

40. If the sequencing cost and time for sufficient coverage becomes prohibitive, a few enrichment strategies can be used. A restriction enzyme-based approach like AlphaHOR-RES relies on preferential digestion of DNA outside of target regions followed by size selection to maintain larger on-target fragments^9,20^. The ONT Cas9 Sequencing Kit (SQC-CS9109) is another option to selectively ligate adapters to targeted regions during library preparation. Adaptive sampling methodologies can also be used with no changes to the wet lab protocol^21,22^.

### Library preparation & sequencing

41. Perform library preparation and sequencing.

CAUTION We have validated the following reagents, but extraction, library preparation, and sequencing reagents are improving rapidly. The important considerations are to choose a DNA extraction method that maintains long DNA molecules, to perform amplification-free library preparation, and to use a flow cell that is compatible with mA calling. These are workflows we have validated and modifications we have made.

#### (A) Target N50 ∼20 kb

TIMING 3 hours

(i) If multiplexing samples on a flow cell, follow Nanopore protocol for Native Barcoding Ligation Kit 1-12 and Native Barcoding Ligation Kit 13-24 with ONT SQK-LSK109. If not multiplexing, use ONT SQK-LSK110. We recommend the following modifications:

a. Load ∼3 µg DNA into end repair.
b. Incubate for 10 minutes at 20°C for end repair instead of 5 minutes.
c. Load ∼ 1 µg of end repaired DNA into barcode ligation.
d. Double the ligation incubation time(s) to at least 20 minutes.
e. Elute in 18 µl instead of 26 µl following barcode ligation reaction cleanup to allow for more material to be loaded into the final ligation.
f. Load ∼3 µg of pooled barcoded material into the final ligation. If needed, concentrate using speedvac to be able to load 3 µg into the final ligation.
g. Perform final elution in 13 µl EB. Take out 1 µl to dilute 1:5 for quantification by Qubit (and size distribution analysis by TapeStation / Bioanalyzer if desired).
h. Load ∼1 µg of DNA onto the sequencer. Input requirements vary by sequencing kit and are becoming lower.

TROUBLESHOOTING

#### (B) Target N50 ∼50 kb

TIMING 5 days

(i) Follow Nanopore protocol for ONT SQK-LSK110 (method validated with this kit only, not with multiplexing with ONT SQK-LSK109) with the following modifications (inspired by Kim et al, dx.doi.org/10.17504/protocols.io.bdfqi3mw^30^):

a. Increase end preparation time to 1 hour with a 30-minute deactivation.
b. Following end preparation, perform a cleanup by combining 60 μL SRE buffer from Circulomics (SS-100-101-01) with the 60 μL end prep reaction.
c. Centrifuge this reaction at 10,000 x g at room temperature for 30 minutes (or until DNA has pelleted).
d. Wash pelleted DNA with 150 μL of 70% ethanol two times, using a 2 minute spin at 10,000 x g between washes.
e. Resuspend the pellet in 31 μL EB.
f. Incubate at 50°C for 1 hour. Incubate at 4°C for at least 48 hours.
g. For the ligation step, reduce ligation volume by half (total of 30 μL DNA in a 50 μL reaction volume). Increase the ligation incubation to 1 hour.
h. Pellet DNA at 10,000 x g at room temperature for 30 minutes.
i. Wash the pellet twice with 100 μL LFB, using a 2 minute spin at 10,000 x g between washes.
j. Resuspend the pellet in 31 μL EB.
k. Incubate at least 48 hours at 4°C.
l. Load 500 ng of DNA onto the sequencer. Input requirements vary by sequencing kit and are becoming lower.
m. If you see the number of active pores has dropped considerably after 24 hours, you can recover pore activity using the flow cell wash kit, then loading additional library material.

TROUBLESHOOTING

### Analysis

42. Perform modified basecalling.
43. (Optional) Use the *dimelo*^25^ Python package for quality control and data visualization. TROUBLESHOOTING

## Timing

### N50 ∼20 kb

Day 1

Steps 1-13, buffer preparation, nuclear permeabilization, primary antibody binding: 2 h

Day 2

Steps 14-39, primary antibody wash, pA binding, activation, and DNA extraction: 8 h (9 h if fixation was performed)

Day 3

Step 41, perform library preparation & start sequencing: 3 h

Day 4

Step 41, re-load sequencer if necessary: 1 h

Day 5

Step 41, re-load sequencer if necessary: 1 h

Day 6

Step 42-43, basecall & perform initial analysis with *dimelo*: 10 h

### N50 ∼50 kb

Day 1

Steps 1-13, buffer preparation, nuclear permeabilization, primary antibody binding: 2 h

Day 2

Steps 14-39, primary antibody wash, pA binding, activation, and DNA extraction: 9 h (10 h if fixation was performed)

Day 3

Step 41, perform library preparation end repair and clean: 2 h

Day 5

Step 41, perform library preparation ligation and clean: 2 h

Day 7

Step 41, start sequencing

Day 8

Step 41, re-load sequencer if necessary: 1 h

Day 9

Step 41, re-load sequencer if necessary: 1 h

Day 10

Step 42-43, basecall & perform initial analysis with *dimelo*: 10 h

## Troubleshooting

Troubleshooting advice can be found in Table 1.

**Table 1.**
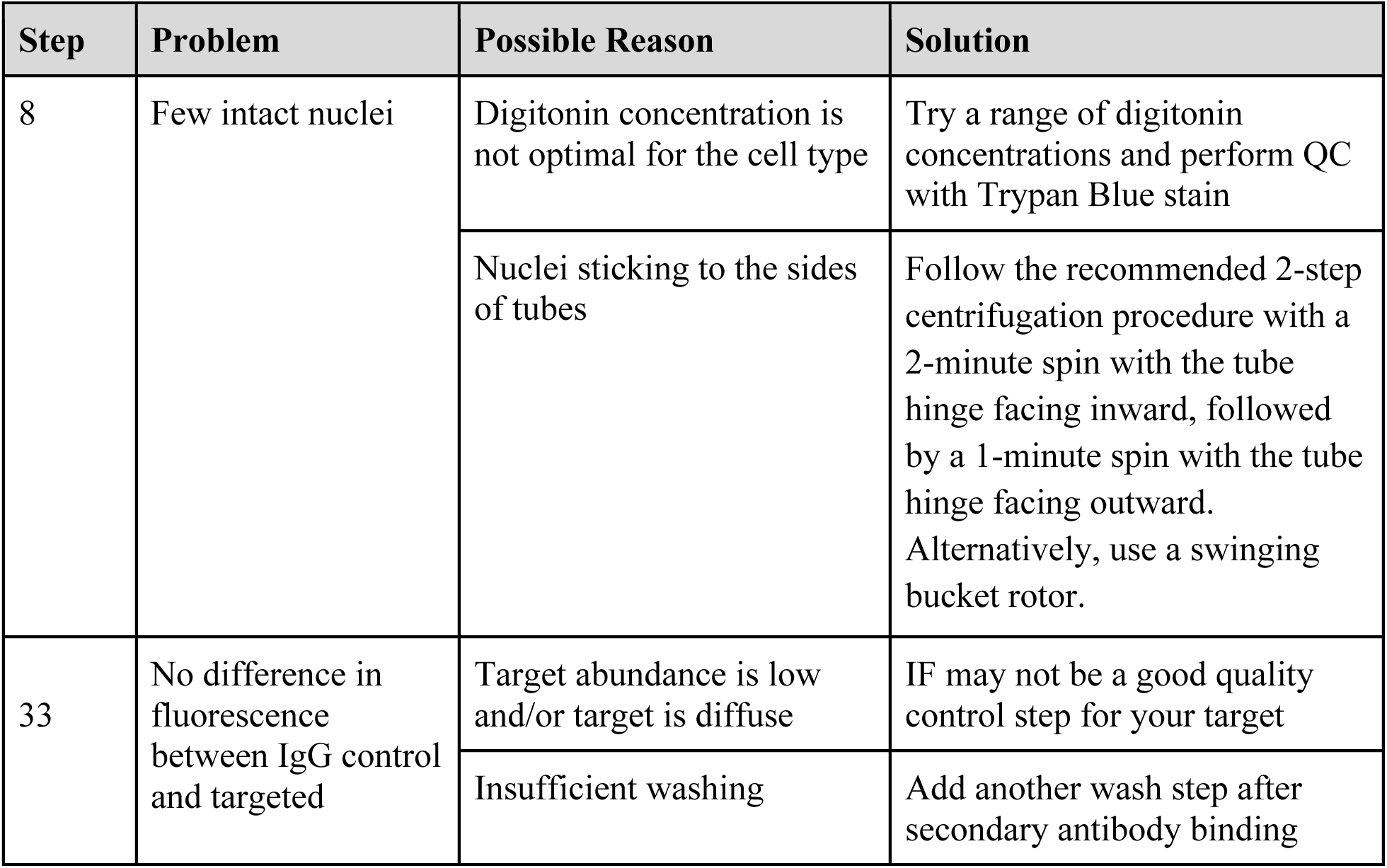
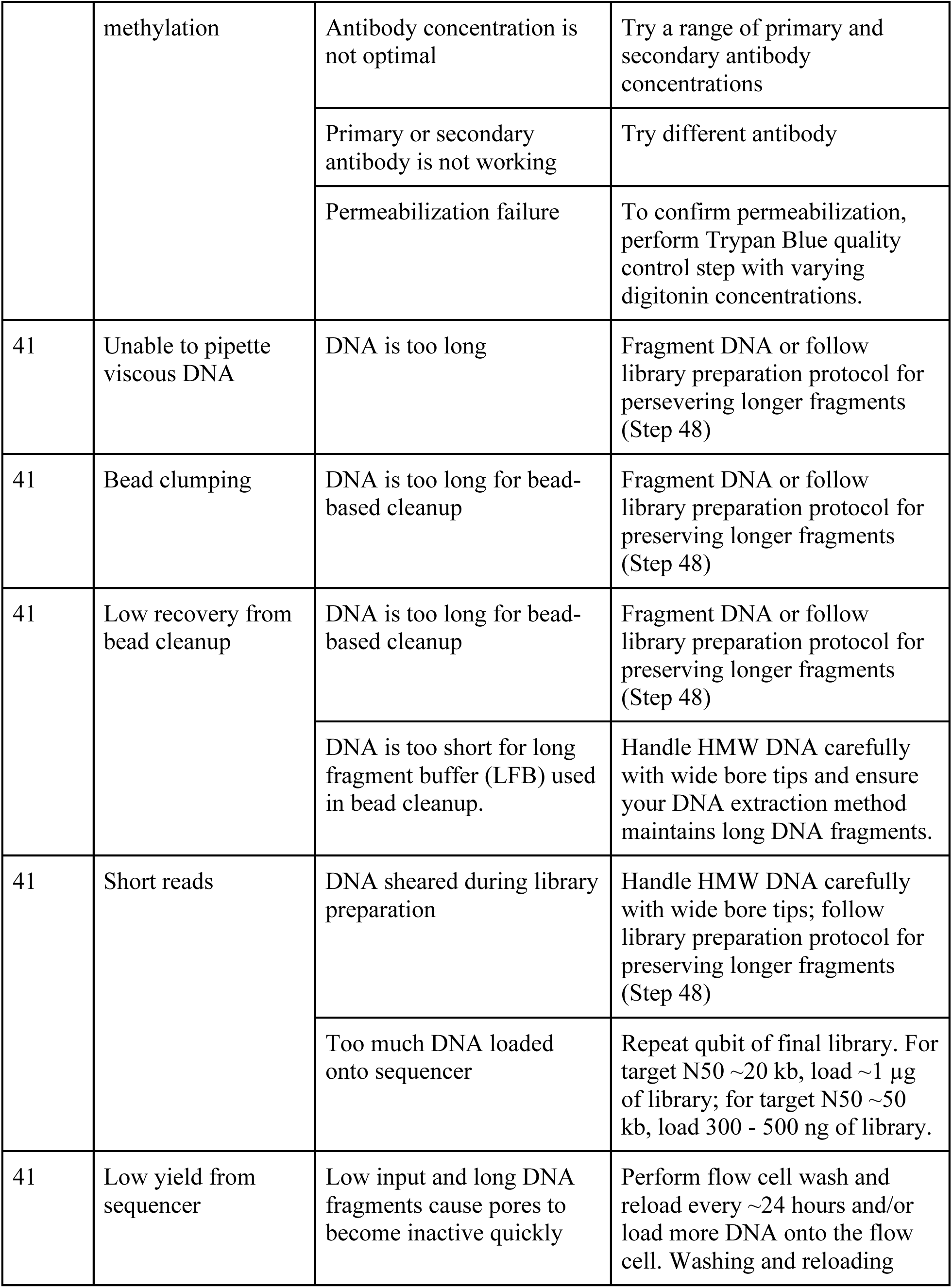
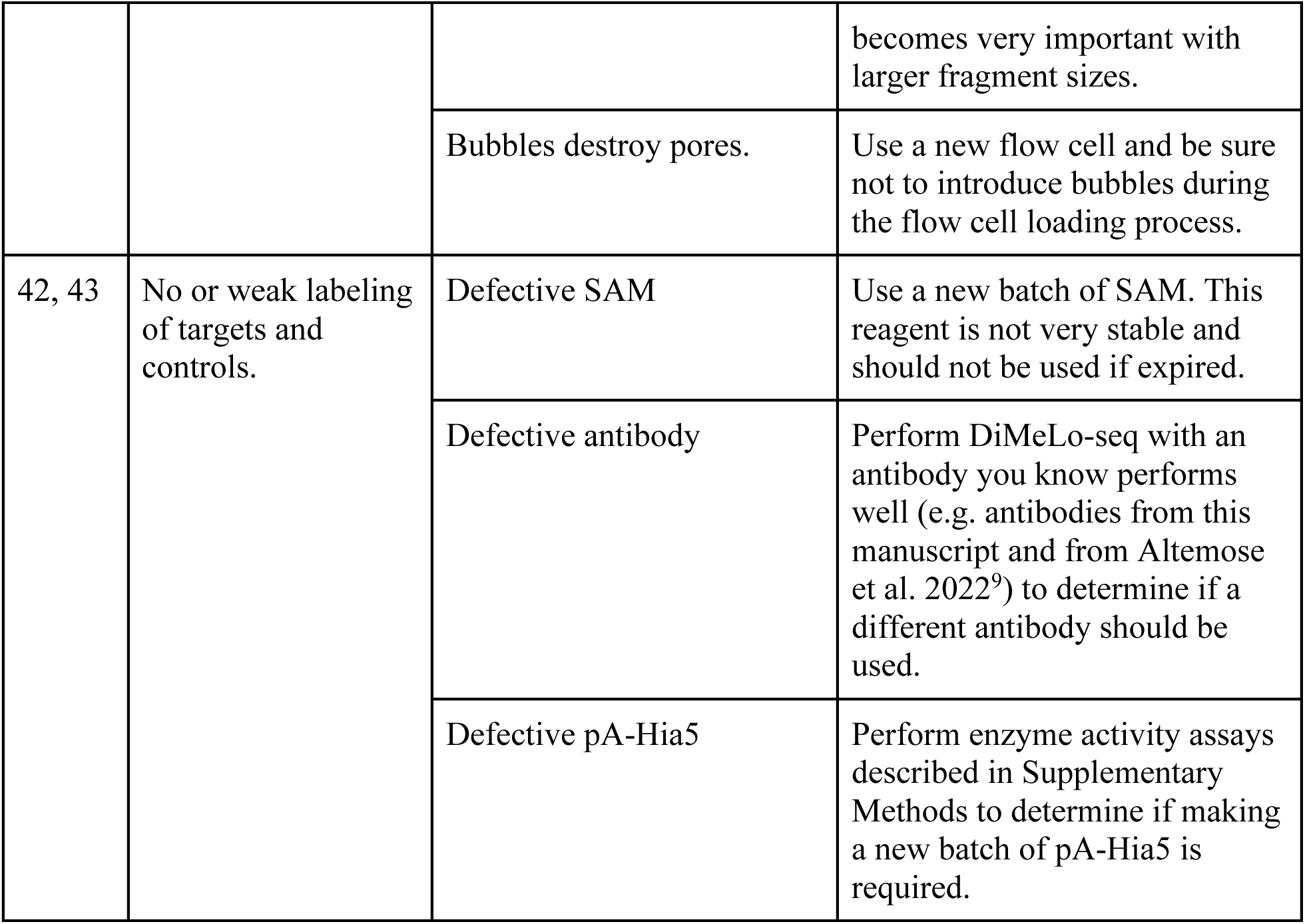
Troubleshooting tips.

## Anticipated results

One person can collect and analyze sequencing data within 3-8 days of beginning the DiMeLo-seq protocol. In this section, we show representative data from DiMeLo-seq experiments targeting H3K27ac, H3K27me3, and H3K4me3 in GM12878 cells and H3K9me3 in *D. melanogaster* embryos (Table 2). We use these targets to provide example output from the *dimelo* package and include suggested figures to evaluate performance and to perform exploratory analysis with DiMeLo-seq data.

**Table 2:** Experimental overview. Summary of experimental specifications for histone modifications profiled using DiMeLo-seq.

| Target | Cell type | Antibody | Library prep kit | Flow cell chemistry | Device | Gb | Cove rage | N50 (bp) |
| --- | --- | --- | --- | --- | --- | --- | --- | --- |
| H3K27ac | GM12878 | Active Motif 39133 | SQK-LSK110 | R9.4.1 | PromethION | 124 | 41X | 25,536 |
| H3K27me3 |  | Active Motif 39055 |  |  |  | 122 | 41X | 27,226 |
| H3K4me3 |  | Active Motif 39916 |  |  |  | 124 | 41X | 25,163 |
| H3K9me3 | *D. melanogaster* embryo | Active Motif 39062 |  |  | MinION | 8.24 | 46X | 27,843 |

The specificity and efficiency of methylation vary by target, depending on the antibody quality, how broad the binding domain is, and the chromatin environment, among other factors. The on-target and off-target methylation levels when targeting H3K27ac, H3K27me3, and H3K4me3 with DiMeLo-seq are shown in Figure 5a-c. These plots are generated from the `plot_enrichment` function. For H3K27ac, to define on-target regions, we used top ChIP-seq peaks for H3K27ac (ENCODE ENCFF218QBO^31^). For off-target, we used top ChIP-seq peaks for H3K27me3 (ENODE ENCFF119CAV^31^). We similarly analyze on- and off-target for H3K27me3 with H3K27me3 top ChIP-seq peaks for on-target and H3K27ac top ChIP-seq peaks for off-target regions. For H3K4me3, to define on-target regions, we used top ChIP-seq peaks for H3K4me3 (ENCODE ENCFF228TWF^31^); for off-target, we used transcription start sites of unexpressed genes where H3K4me3 is not expected to be present. The on-target methylation levels are higher for H3K27me3 compared to H3K27ac, despite H3K27me3 being a repressive mark in a less accessible genomic context. This is likely because it binds a broader genomic region, allowing a larger methylated footprint. The performance difference could also occur if the anti-H3K27me3 antibody performs better than the anti-H3K27ac antibody used in these experiments. The off-target methylation level is also higher in H3K27me3 compared to H3K27ac. This is likely because the off-target region used in this analysis is H3K27ac ChIP-seq peaks, which are in very accessible regions of the genome, and off-target methylation with DiMeLo-seq occurs preferentially within open chromatin. Again, higher off-target methylation can also be caused by differences in antibody performance.

**Figure 5.**
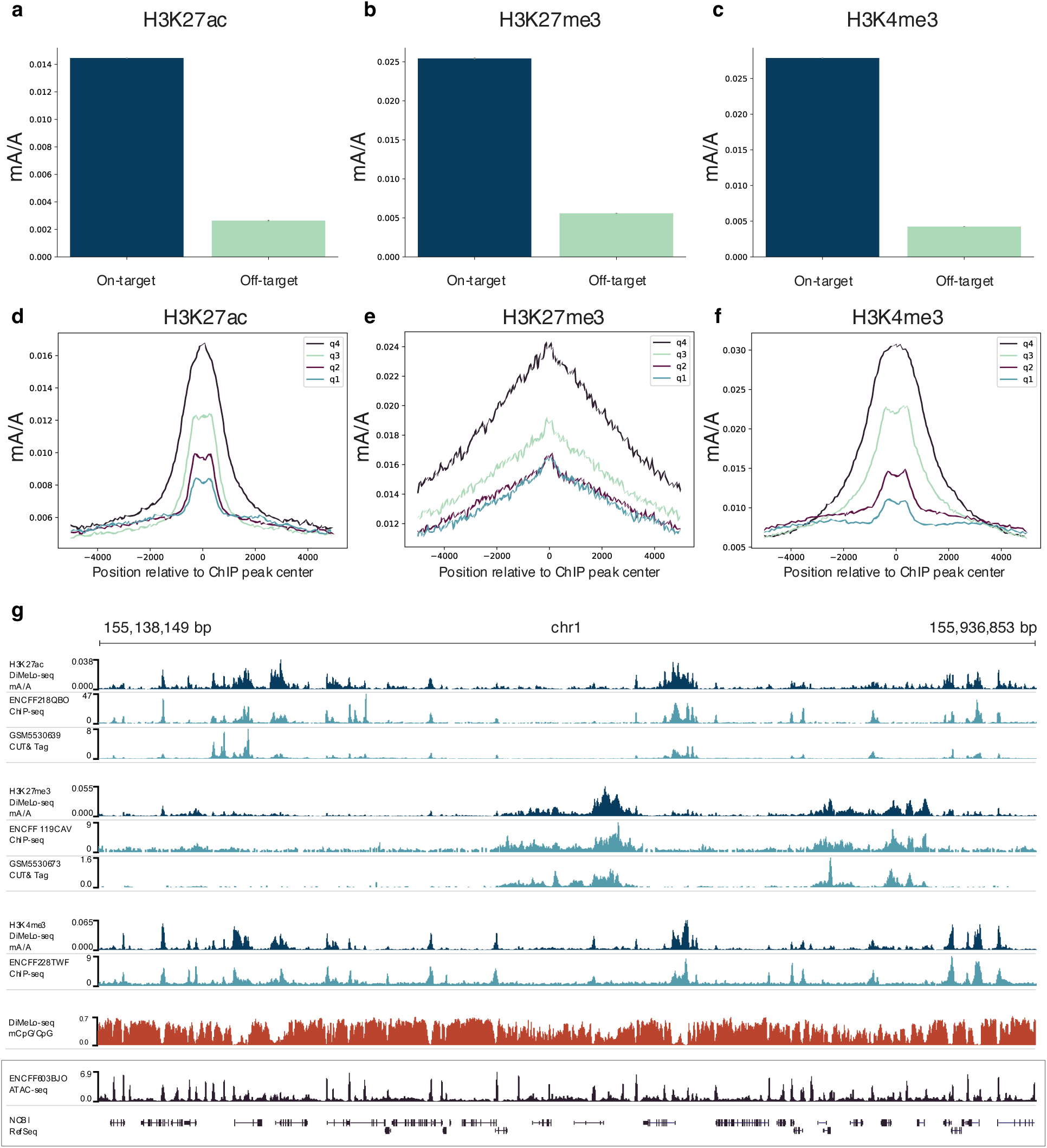
Validation of targeted methylation. **a-c,** Using BED files defining on- and off-target regions, the `plot_enrichment` function can be used to determine whether methylation is concentrated within expected regions. We’ve defined on-target regions using ChIP-seq peaks for the corresponding histone marks. We defined off-target regions when targeting H3K27ac as H3K27me3 ChIP-seq peaks and when targeting H3K27me3 as H3K27ac ChIP-seq peaks; for off-target regions for H3K4me3 we use transcription start sites for unexpressed genes. Methylation probability threshold of 0.75 was used. Error bars represent 95% credible intervals determined for each ratio by sampling from posterior beta distributions computed with uninformative priors. **d-f,** Methylation profiles centered at ChIP-seq peaks for H3K27ac-, H3K27me3-, and H3K4me3-targeted DiMeLo-seq are plotted using `plot_enrichment_profile`. The quartiles (q4 to q1) indicate the strength of the ChIP-seq peaks which the DiMeLo-seq reads overlap. Methylation probability threshold of 0.75 was used. **g,** Aggregate browser traces comparing DiMeLo-seq signal to ChIP-seq and CUT&Tag. BED files used for creating aggregate curves are generated either from `parse_bam` or `plot_browser`. CpG methylation signal is aggregated from the H3K27ac-, H3K27me3-, and H3K4me3-targeted DiMeLo-seq experiments. Methylation probability threshold of 0.8 was used. ATAC-seq and NCBI RefSeq annotations are also shown.

The methylation profile centered at features of interest can be visualized using the `plot_enrichment_profile` function. Here, we show profiles from H3K27ac-, H3K27me3-, and H3K4me3-targeted DiMeLo-seq with aggregate methylation curves from reads centered at ChIP-seq peaks of varying strength (Figure 5d-f) (ENCODE ENCFF218QBO, ENCFF119CAV, ENCFF228TWF^31^). H3K27ac and H3K4me3 have narrow peaks, while H3K27me3 has a broader peak. Signals for all three marks track with ChIP-seq peak strength, indicating concordance between DiMeLo-seq and ChIP-seq in aggregate.

To qualitatively demonstrate the concordance between DiMeLo-seq and other methods for measuring protein-DNA interactions - here ChIP-seq and CUT&Tag - we created aggregate browser tracks across a stretch of chromosome 1 (Figure 5g) (ENCODE ENCFF218QBO, ENCFF119CAV, ENCFF228TWF; GEO GSM5530639, GSM5530673^31,32^). DiMeLo-seq signal for all three histone marks tracks with ChIP-seq and CUT&Tag profiles. These curves were generated using the BED file output from the `plot_browser` function with smoothing in a 100 bp window. DiMeLo-seq also measures endogenous CpG methylation together with protein binding. An aggregate mCpG signal from the three DiMeLo-seq samples is shown, and dips in mCpG are evident where H3K27ac and H3K4me3 signals are highest. H3K27ac and H3K4me3 are both marks of open chromatin and have peaks overlapping accumulations in ATAC-seq signal (ENCODE ENCFF603BJO^31^).

In addition to comparing DiMeLo-seq to other methods, we also evaluated methylation profiles around genomic features where our targets are expected to localize. In particular, both H3K27ac and H3K4me3 are found at transcription start sites (TSSs)^33^. Using the `plot_enrichment_profile` function, we created the aggregate methylation and single-molecule methylation plots shown in Figure 6a. As expected, both marks have enrichment at the TSSs, with the highest methylation levels at the TSSs for the genes with highest expression.^33^ The periodicity from positions 0 bp to 500-1000 bp with respect to the TSS indicate preferential methylation of linker DNA between strongly positioned nucleosomes downstream from the TSSs for both targets. For genes that are not expressed (quartile 1), no significant enrichment at TSSs is evident.

**Figure 6.**
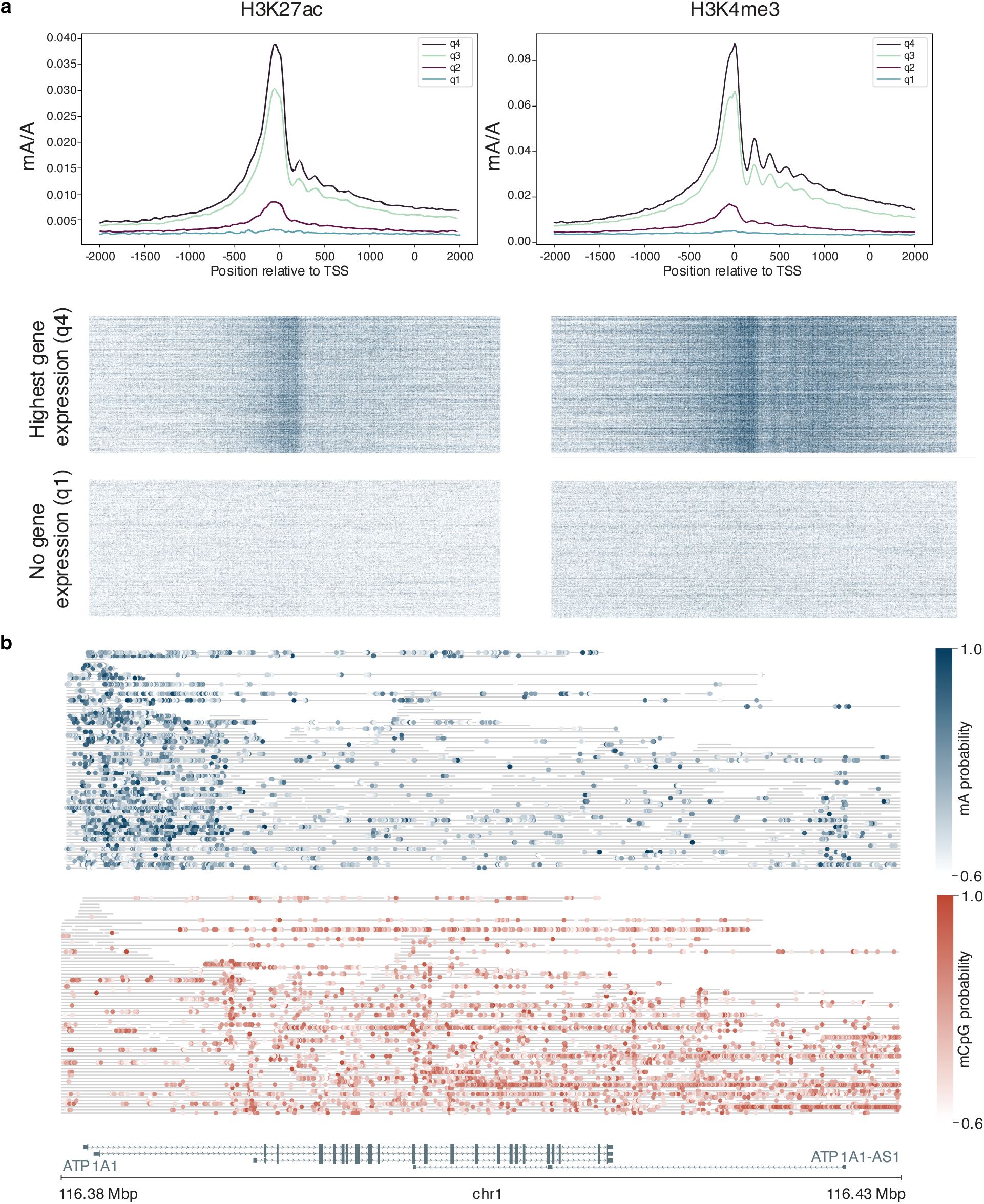
Evaluating protein binding at regions of interest. Both H3K27ac and H3K4me3 are found at transcription start sites (TSS). **a,** Signal from H3K27ac- and H3K4me3-targeted DiMeLo-seq at TSS. Reads overlapping TSS, gated by gene expression level from highest gene expression (quartile 4 (q4)) to lowest gene expression (quartile 1 (q1)). Aggregate mA/A profiles are shown for all reads spanning these TSSs. Single molecules are shown below with blue representing mA calls for TSS for the highest gene expression (q4) and for no gene expression (q1). Aggregate and single-molecule plots were produced with `plot_enrichment_profile`. Methylation probability threshold of 0.75 was used. **b,** Single-molecule browser plots produced from `plot_browser` from H3K4me3-targeted DiMeLo-seq experiment. Using a methylation probability threshold of 0.6, mA (top, blue) and mCpG (bottom, red) calls are shown for the same molecules (grey lines). NCBI RefSeq genes are shown below.

Using the `plot_browser` function, single molecules are shown from H3K4me3-targeted DiMeLo-seq in a window around a highly expressed gene in GM12878 (Figure 6b). Methylated adenines are enriched around the TSS for the highly expressed gene ATP1A1. Together with mA, the endogenous mCpG can also be analyzed. Here, it is evident that mCpG is depleted in the regions around TSS where H3K4me3 is enriched, as has been previously reported^34^. Multiple TSSs are spanned by some of the molecules in the region from 116.38 Mbp to 116.43 Mbp on chromosome 1, highlighting DiMeLo-seq’s ability to probe multiple binding events on a single molecule.

DiMeLo-seq can be used to target proteins in nuclei not only from cultured cells but also from primary tissue or intact organisms. We mapped H3K9me3 distributions in *D. melanogaster* embryos across the genome and show that averaging methylation signal from single molecules generates profiles consistent with previously published ChIP-seq data (Figure 7)^35^. DiMeLo-seq coverage is consistent across the entire *D. melanogaster* genome because DiMeLo-seq’s long reads can be mapped in repetitive regions of the genome. Spikes in coverage that also appear as spikes in DiMeLo-seq and ChIP-seq signal are due to centromeric repetitive sequences that are missing from the reference. This same repetitive DNA is where H3K9me3 is abundant, resulting in corresponding spikes in DiMeLo-seq methylation signal as well. We highlight a transition on chr3L where H3K9me3 accumulates and show that the accumulation is evident on single molecules using the `plot_browser` function.

**Figure 7.**
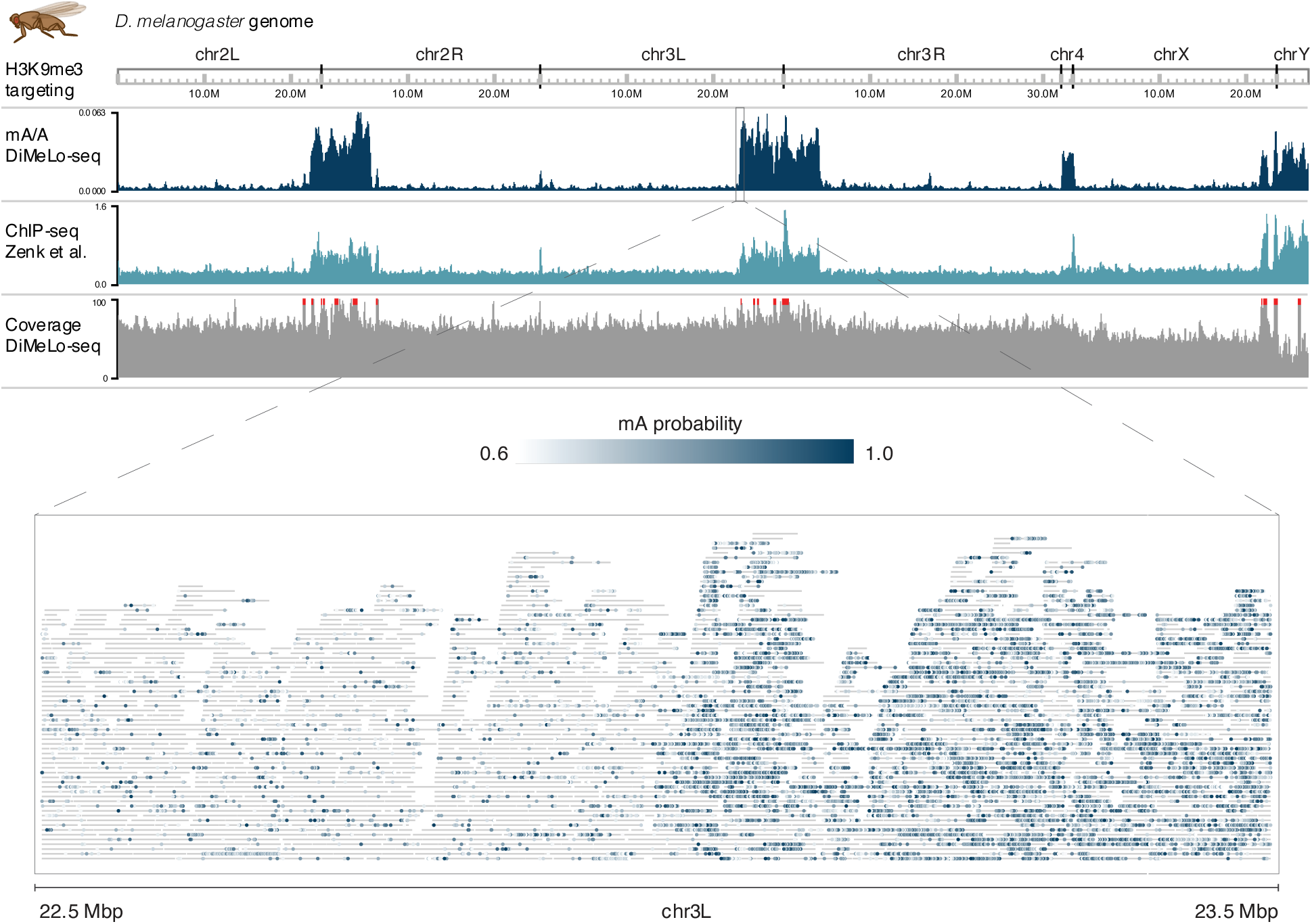
H3K9me3-targeted DiMeLo-seq in *D. melanogaster* embryos. Aggregate mA/A across the entire *D. melanogaster* genome from a DiMeLo-seq experiment targeting H3K9me3 is shown in dark blue. H3K9me3 ChIP-seq data in *D. melanogaster* embryos is shown in light blue.^35^ Coverage from the DiMeLo-seq experiment is shown in grey. A region on chr3L where a transition from H3K9me3 depletion to H3K9me3 enrichment is highlighted with a single-molecule browser plot generated from `plot_browser`. Grey lines indicate reads and blue dots indicate mA calls with intensity colored by probability of methylation. An alignment length filter of 10 kb was applied. Methylation probability threshold of 0.6 was used.

It is important to note that the fraction of bases reported as modified should not be interpreted as the fraction of cells or molecules that have protein bound. First, this fraction is averaged across all adenines on all reads overlapping a given window and thus is an aggregate rather than a single-molecule statistic. Second, this fraction varies considerably with the modification probability threshold used. Third, the bin size used for creating aggregate plots influences the computed fractions. To measure the single-molecule interaction frequency, analysis should be performed as described for LMNB1 in Altemose et al.^9^

The DiMeLo-seq protocol described here enables profiling of protein-DNA interactions in repetitive regions of the genome, makes phasing easier for determining haplotype-specific interactions^9^, detects joint binding events on single molecules of DNA, and captures protein binding together with endogenous CpG methylation. These analyses are uniquely enabled by the use of long-read, native sequencing because short, clonally amplified reads cannot map to repetitive regions, span fewer SNPs making phasing more difficult, cannot store information about multiple binding events on a single read, and lose endogenous CpG methylation information through amplification. DiMeLo-seq performance varies by protein target, antibody quality, and chromatin environment; therefore, methylation sensitivity and specificity must be evaluated for each new target. The *dimelo* software package provides tools for quality control and data exploration for the multimodal datasets that DiMeLo-seq produces.

## Data availability

Raw sequencing data are available in the Sequence Read Archive (SRA) under BioProject accession PRJNA855257 and processed data are available on Gene Expression Omnibus (GEO) under accession GSE208125 (https://www.ncbi.nlm.nih.gov/geo/query/acc.cgi?acc=GSE208125). All raw fast5 sequencing data from the accompanying Altemose et al. manuscript are available in the Sequence Read Archive (SRA) under BioProject accession PRJNA752170. H3K27ac ChIP-seq data in GM12878 available from ENCODE Project Consortium under accession ENCFF218QBO (https://www.encodeproject.org/files/ENCFF218QBO/). H3K27me3 ChIP-seq data in GM12878 available from ENCODE Project Consortium under accession ENCFF119CAV (https://www.encodeproject.org/files/ENCFF119CAV/). H3K4me3 ChIP-seq data in GM12878 available from ENCODE Project Consortium under accession ENCFF228TWF (https://www.encodeproject.org/files/ENCFF228TWF/). H3K27ac CUT&Tag data in GM12878 available on Gene Expression Omnibus (GEO) under accession GSM5530639 (https://www.ncbi.nlm.nih.gov/geo/query/acc.cgi?acc=GSM5530639). H3K27me3 CUT&Tag data in GM12878 available on GEO under accession GSM5530673 (https://www.ncbi.nlm.nih.gov/geo/query/acc.cgi?acc=GSM5530673).

ATAC-seq data in GM12878 available from ENCODE Project Consortium under accession ENCFF603BJO (https://www.encodeproject.org/files/ENCFF603BJO/). Transcription start site and gene annotations from NCBI RefSeq downloaded from UCSC Genome Browser (https://genome.ucsc.edu/cgi-bin/hgTrackUi?g=refSeqComposite&db=hg38). RNA-seq data in GM12878 available from ENCODE Project Consortium under accession ENCFF978HIY (https://www.encodeproject.org/files/ENCFF978HIY/). *D. melanogaster* H3K9me3 ChIP-seq data available on GEO under accession GSE140539 (https://www.ncbi.nlm.nih.gov/geo/query/acc.cgi?acc=GSE140539). File GSE140539_H3K9me3_sorted_deepnorm_log2_smooth.bw was used.

## Code availability

The *dimelo*^25^ Python package for analysis of DiMeLo-seq data is available on Github: https://github.com/streetslab/dimelo

## Author contributions

AM, NA, and AS designed the study. AM, NA, and LDB performed the experiments. AM, RM, and JM developed *dimelo* software package. AM and NA analyzed and interpreted the data. AM, NA, and RM made the figures. AM wrote the manuscript, with input from NA, RM, JM, LDB, KS, GK, AFS, and AS. AS and NA supervised the study.

## Acknowledgements

This work was supported by the Chan Zuckerberg Biohub and by the NIGMS of the National Institutes of Health under award number R01HG012383 to AS, R01GM074728 to AFS, and R35GM139653 to GHK. AM is supported by a NSF GRFP award. NA is an HHMI Hanna H. Gray Fellow. AS is a Chan Zuckerberg Biohub Investigator, and a Pew Scholar in the Biomedical Sciences. The sequencing was carried out by the DNA Technologies and Expression Analysis Core at the UC Davis Genome Center, supported by NIH Shared Instrumentation Grant 1S10OD010786-01. This project has been made possible in part by grant number 2022-253563 from the Chan Zuckerberg Initiative DAF, an advised fund of Silicon Valley Community Foundation.

## Supplementary Methods

### Additional protocol and material availability

DiMeLo-seq: https://doi.org/10.17504/protocols.io.b2u8qezw (version 2 of https://www.protocols.io/view/dimelo-seq-directed-methylation-with-long-read-seq-n2bvjxe4wlk5 used for this manuscript); pA–Hia5 protein purification: https://doi.org/10.17504/protocols.io.bv82n9ye (version 1 of https://www.protocols.io/view/pa-hia5-protein-expression-and-purification-x54v9j56mg3e used for this manuscript); AlphaHOR-RES: https://doi.org/10.17504/protocols.io.bv9vn966 (version 1 of https://www.protocols.io/view/alphahor-res-a-method-for-enriching-centromeric-dn-ewov141rovr2 used for this manuscript). Plasmids are available on Addgene: pA–Hia5 expression plasmid (pET-pA–Hia5; Addgene, 174372) and pAG–Hia5 expression plasmid (pET-pAG–Hia5; Addgene, 174373).

### Cell culture

GM12878 cells (GM12878, Coriell Institute; mycoplasma tested; https://scicrunch.org/resolver/RRID:CVCL_7526) were maintained in RPMI-1640 with L-glutamine (Gibco, 11875093) supplemented with 15% FBS (VWR 89510-186) and 1% penicillin–streptomycin (Gibco, 15070063) at 37 °C in 5% CO2.

#### Antibody information

Antibodies used in this study were: (1) Histone H3K27ac antibody (pAb) (Active Motif 39133 https://scicrunch.org/resolver/RRID:AB_2722569), (2) Histone H3K27me3 antibody (pAb) (Active Motif 39055, https://scicrunch.org/resolver/RRID:AB_2561020), (3) Histone H3K4me3 antibody (pAb) (Active Motif 39916, https://scicrunch.org/resolver/RRID:AB_2687512), (4) Histone H3K9me3 antibody (pAb) (Active Motif 39062, https://scicrunch.org/resolver/RRID:AB_2532132).

### DiMeLo-seq in GM12878 cells

For each target, 3.24 M cells from fresh culture were input to DiMeLo-seq. Antibody dilutions were all 1:50. DNA extraction was performed using the Monarch Genomic DNA Purification Kit (NEB T3010S).

### DiMeLo-seq analysis

Basecalling was performed using Oxford Nanopore Technologies’ Megalodon software version 2.3.1 and Guppy version 4.5.4 with the Rerio res_dna_r941_min_modbases-all-context_v001.cfg basecalling model. We used the *dimelo*^25^ Python package (https://github.com/streetslab/dimelo) for plotting.

### *DiMeLo-seq with* D. melanogaster *embryos*

#### Timed embryo collections for downstream DiMeLo-seq

Approximately 200-400 OregonR flies were maintained on standard molasses medium before transfer to embryo collection cages with apple-juice plates and yeast paste. A heterogeneous mixture of embryos were collected overnight from 2-4 cages and pooled together. Embryos were rinsed from the apple-juice plates with DI water and collected in a mesh sieve, the chorion was removed by soaking in 50% bleach for 90 seconds, and then rinsed with water to remove the bleach. Embryos were transferred to a 1.5 mL cryotube, allowed to settle, and the water was replaced with 1 mL Embryo Storage Buffer. Embryos were frozen in a Mr. Frosty isopropanol bath at -80°C overnight, then stored at -80°C. Nuclei were prepped for DiMeLo-seq by thawing the embryos at room temperature, removing the storage buffer and replacing it with 1mL of 1xPBS. Embryos were transferred to a 1 mL glass Dounce homogenizer and lysed with 10-15 strokes of a loose-fitting pestle. Nuclei were gently pelleted at 600 x g for 3 minutes at 4°C, the supernatant was removed, and the pellet was resuspended in Dig-wash buffer for downstream DiMeLo-seq. DNA extraction was performed using the Monarch HMW DNA Extraction Kit (NEB T3050L) with 2000 rpm at lysis.

#### Materials

Apple juice plates

Yeast paste

Oregon R flies (young) ∼ 200 per bottle

Bleach

DI water

#### Buffers

Embryo Storage Buffer: 80% Schneider’s S2 media (Thermo 21720024), 10% FBS, 10% DMSO 1X PBS

50% bleach solution

#### Equipment

Small embryo sieve

Paintbrush

Squirt bottle (with DI water)

1 mL pipette and tips

Mr. Frosty isopropanol freezing bath

1.5 mL Eppendorf tubes

### Library preparation and sequencing

All library preparation was performed using ONT SQK-LSK110 with the standard protocol’s bead-based cleanup protocol. Targets in GM12878 were sequenced on PromethION with R9.4.1 flow cells (ONT FLO-PRO002). *D. melanogaster* embryo experiments were sequenced on MinION with R9.4.1 flow cells (ONT FLO-MIN106D).

### Measuring pA-Hia5 methyltransferase activity

Enzyme activity can vary by batch, so we have used a few approaches to assay methyltransferase activity. This benchmarking is important for quantitative comparisons across samples. It is advisable to use the same pA-Hia5 batch across conditions for comparative analyses. Some options for benchmarking activity include:

(1) Perform methylation-sensitive restriction enzyme digestion following incubation of a known DNA sequence that has GATC site(s) with pA-Hia5 in activation buffer. Run the digest on a gel or on the TapeStation. Use either DpnI or DpnII. DpnI will cut only at GmATC and DpnII will cut only at GATC without methylation. This restriction-enzyme-based strategy was demonstrated in Altemose et al.^9^ A user can also order the below oligos, anneal them, methylate with pA-Hia5, perform methylation-sensitive restriction enzyme digestion, and then run the TapeStation or a gel.jhgjhg

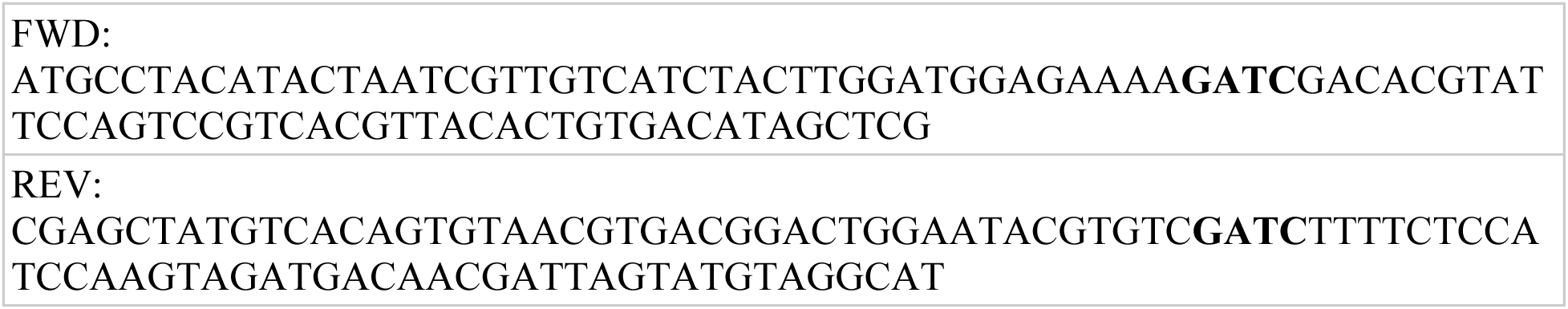
(2) Perform dot blots as described by Stergachis et al.^11^
(3) For users performing many DiMeLo-seq experiments, another method for validating enzymatic activity is to perform an additional DiMeLo-seq experiment in parallel, using a known antibody and well characterized target and cell type. This provides a positive control for performance assessment from batch to batch. For example, we have published expected methylation levels in on-target and off-target regions when targeting LMNB1 in HEK293T cells^9^.

## Supplementary Material

**Supplementary Table 1.** Comparison of protein A and protein A/G performance. We targeted LMNB1 in HEK293T cells and methylated with either pA-Hia5 or pA/G-Hia5 from within one experimental batch. On-target methylation was defined as the fraction of adenines methylated in constitutive lamina associated domains. Off-target methylation was defined as the fraction of adenines methylated in constitutive inter lamina associated domains. A basecalling probability threshold of 0.8 was used for calling methylation.

| protein | On-target / off-target methylation | On-target methylation (mA/A) |
| --- | --- | --- |
| pA-Hia5 | 19.4698 | 0.0079 |
| pA-Hia5 | 17.5034 | 0.0059 |
| pA/G-Hia5 | 19.3952 | 0.0046 |
| pA/G-Hia5 | 18.6125 | 0.0042 |

